# A single-cell RNA expression map of human coronavirus entry factors

**DOI:** 10.1101/2020.05.08.084806

**Authors:** Manvendra Singh, Vikas Bansal, Cédric Feschotte

## Abstract

To predict the tropism of human coronaviruses, we profile 28 SARS-CoV-2 and coronavirus-associated receptors and factors (SCARFs) using single-cell RNA-sequencing data from a wide range of healthy human tissues. SCARFs include cellular factors both facilitating and restricting viral entry. Among adult organs, enterocytes and goblet cells of the small intestine and colon, kidney proximal tubule cells, and gallbladder basal cells appear most permissive to SARS-CoV-2, consistent with clinical data. Our analysis also suggests alternate entry paths for SARS-CoV-2 infection of the lung, central nervous system, and heart. We predict spermatogonial cells and prostate endocrine cells, but not ovarian cells, to be highly permissive to SARS-CoV-2, suggesting male-specific vulnerabilities. Early stages of embryonic and placental development show a moderate risk of infection. The nasal epithelium looks like another battleground, characterized by high expression of both promoting and restricting factors and a potential age-dependent shift in SCARF expression. Lastly, SCARF expression appears broadly conserved across human, chimpanzee and macaque organs examined. Our study establishes an important resource for investigations of coronavirus biology and pathology.

## INTRODUCTION

The zoonotic spillover of the novel severe acute respiratory syndrome coronavirus 2 (SARS-CoV-2) in the human population is causing a disease known as coronavirus disease 2019 (COVID-19) (Lu et al., 2020; Paules et al., 2020). Since the first case reported in late December 2019, SARS-CoV-2 has spread to 203 countries, infecting more than 4 million humans and claiming over 300,000 lives, primarily among the elderly (John Hopkins University and Medicine, 2020; Xu and Li, 2020; Zhou et al., 2020). SARS-CoV-2 is the third coronavirus, after SARS-CoV and MERS-CoV, causing severe pneumonia in humans (Corman et al., 2018). Of these, SARS-CoV-2 is most closely related to SARS-CoV in nucleotide sequence (∼80%), but all three coronaviruses appear to cause similar pathologies (Ding et al., 2003; Lu et al., 2020; Ng et al., 2016).

Emerging clinical and molecular biology data from COVID-19 patients have detected SARS-CoV-2 nucleic acids primarily in bronchoalveolar lavage fluid, sputum, and nasal swabs, and less frequently in fibrobronchoscope brush biopsies, pharyngeal swabs, feces and with even lower positive rates in blood and urine (Ling et al., 2020; Wang et al., 2020d; Wu et al., 2020a; Young et al., 2020; Zou et al., 2020a). Pathological investigations, including post-mortem biopsies, remain limited for COVID-19, but have confirmed major pulmonary damage as the most likely cause of death in the cases examined (Huang et al., 2020; Xu et al., 2020b). There is also growing evidence that SARS-CoV-2 infection can damage other organ systems including the heart, kidney, liver, and gastrointestinal tract, as documented previously for SARS and MERS (Ding et al., 2003; Gu et al., 2005; Ng et al., 2016). Notably, it has been reported that cardiac injury is common in COVID-19 patients (Shi et al., 2020), as well as acute kidney injury (Cheng et al., 2020; Diao et al., 2020; Fanelli et al., 2020; Volunteers et al., 2020; Wang et al., 2020c). Severe COVID-19 patients show frequent liver dysfunctions (Zhang et al., 2020) and evidence of gastrointestinal infection has also been reported (Gao et al., 2020; Xiao et al., 2020). Evidence of impaired gonadal function in male COVID-19 patients was also recently presented (Ma et al., 2020; Wang et al., 2020c). Intriguingly, SARS-CoV-2 can also be detected in the brain or cerebrospinal fluid and may even cause neurological complications (Moriguchi et al., 2020; Wu et al., 2020b). It is not yet understood what causes the wide range of clinical phenotypes observed in people infected with SARS-CoV-2. Importantly, it remains unclear which of these pathologies are caused by direct infection of the organs affected or indirect effects mediated by systemic inflammatory responses or comorbidities. A prerequisite to resolve these critical questions is to gain a better understanding of the tropism of the virus (which tissues and cell types are permissive to SARS-CoV-2 infection) and of the cellular processes and genetic factors modulating the course and outcome of an infection.

Because SARS-CoV-2 is a novel virus, our current knowledge of cellular factors regulating its entry into cells is mostly derived from studies of SARS-CoV, MERS-CoV, and ‘commensal’ human coronaviruses (hCoV). The canonical entry mechanism of these coronaviruses is a two-step process mediated by the viral Spike (S) protein decorating the virion. First, the S protein must bind directly to a cell surface receptor and second, it must be cleaved (‘primed’) by a cellular protease to enable membrane fusion. Thus, the cellular tropism of a coronavirus is conditioned not only by the expression of an adequate receptor on the cell surface, but also by the presence of a host-encoded protease capable of cleaving the S protein, preferably at or close to the site of receptor binding (de Haan and Rottier, 2005; Tang et al., 2020). For both SARS-CoV and SARS-CoV-2, angiotensin-Converting Enzyme 2 (ACE2) and Transmembrane Serine Protease 2 (TMPRSS2) have been identified as a prime receptor and a critical protease, respectively, for entry into a target cell (Glowacka et al., 2011; Hoffmann et al., 2020; Li et al., 2003; Matsuyama et al., 2010; Wrapp et al., 2020). These findings have prompted numerous efforts to profile the basal expression levels of *ACE2* and/or *TMPRSS2* across healthy human tissues in order to predict the tropism of these two closely related viruses. While studies monitoring protein abundance *in situ* (e.g. immunocytochemistry) offer a more direct assessment, and have been conducted previously to study ACE2 and/or TMPRSS2 expression (Hamming et al., 2004; Hikmet et al., 2020; Hoffmann et al., 2020), most recent investigations have taken advantage of single-cell RNA-sequencing (scRNA-seq) data to profile the expression of these two factors at cellular resolution in a wide array of tissues (see references in Table S1).

Collectively these studies have revealed a subset of tissues and cell types potentially susceptible to SARS-CoV-2 (see Table S1 for a summary). However, they suffer from several limitations. First, most studies (15/27) profiled a single organ or organ system, and the majority focused on the respiratory tract. Second, most studies (19/27) restricted their analysis to *ACE2* and/or *TMPRSS2*, ignoring other factors potentially limiting SARS-CoV-2 entry or replication. Yet, there is evidence that these two proteins alone cannot solely explain all the current clinical and research observations. For instance, certain cell lines (e.g. A549 alveolar lung carcinoma) can be infected by SARS-CoV-2 in the absence of appreciable level of *ACE2* RNA or protein (Blanco-Melo et al., 2020; Wyler et al., 2020). Similarly, clinical data point to SARS-CoV-2 infection of several organs, such as lung, bronchus, nasopharynx, esophagus, liver and stomach, where *ACE2* expression could not be detected in healthy individuals (Hikmet et al., 2020; Zou et al., 2020b). Moreover, there are discordant reports as to where and how much *ACE2* may be expressed in certain cells, including alveolar type II cells of the lung, which are widely regarded as a primary site of infection and tissue damage. Together, these observations suggest that either *ACE2* expression levels vary greatly between individuals or during the course of an infection (Ziegler et al., 2020) or that SARS-CoV-2 can use alternate receptor(s) to enter certain cell types. For instance, cell surface protein Basignin (BSG, also known as CD147) has been shown to interact with the S protein in vitro and facilitate entry of SARS-CoV and SARS-CoV-2 in Vero and 293T cells (Vankadari and Wilce, 2020; Wang et al., 2020b). In fact, SARS-CoV and other hCoVs can utilize multiple cell surface molecules to promote their entry into cells, including ANPEP (Yeager et al., 1992), CD209 (DC-SIGN) (Yang et al., 2004), CLEC4G (LSECtin) (Marzi et al., 2004), and CLEC4M (LSIGN/CD299) (Gramberg et al., 2005). Likewise, hCoVs can use a variety of cellular proteases to prime their S protein, in substitution for TMPRSS2 in a cell type-specific manner. These include other members of the TMPRSS family (e.g. TMPRSS4) (Glowacka et al., 2011; Zang et al., 2020), but also Cathepsins (CTSL/M) (Simmons et al., 2013a) and FURIN (Mille and Whittaker, 2014; Walls et al., 2020). To our knowledge, no single study has examined systematically the expression of these alternate hCoV entry factors. Just as importantly, none of the previous studies have taken into account the expression of host factors known to oppose or restrict cellular entry of hCoVs, including SARS-CoV-2, such as LY6E (Pfaender et al., 2020) and IFITM proteins (Huang et al., 2011). Overall, our understanding of cellular factors underlying the potential tropism of SARS-CoV-2 remain very partial.

To begin addressing these gaps, we curated a list of 28 human genes referred to as SCARFs for <u>S</u>ARS-CoV-2 and <u>C</u>oronavirus-<u>A</u>ssociated <u>R</u>eceptors and <u>F</u>actors (Figure 1A and Table S2) and surveyed their basal RNA expression levels across a wide range of healthy tissues. Specifically, we mined publicly available scRNA-seq datasets using consistent normalization procedures to integrate and compare the dynamics of SCARF expression in human pre-implantation embryos (Yan et al., 2013), at the maternal-fetal interface (Vento-Tormo et al., 2018), in male and female gonads (Sohni et al., 2019; Wagner et al., 2020) and 14 other adult tissues (Han et al., 2020), as well as nasal brushing from young and old healthy donors (Deprez et al., 2019; Garcıá et al., 2019; Vieira Braga et al., 2019). Additionally, we use bulk transcriptomics for four organs of interest (lung, kidney, liver, heart) from human, chimpanzee, and macaque (Blake et al., 2020) to examine the conservation of SCARF expression across primates. This study represents the most systematic survey of SCARF expression to date and provides a valuable resource, including a user-friendly web browser, for interpreting and prioritizing clinical, pathological, and biological studies of SARS-CoV-2 and COVID-19.

**Figure 1:**
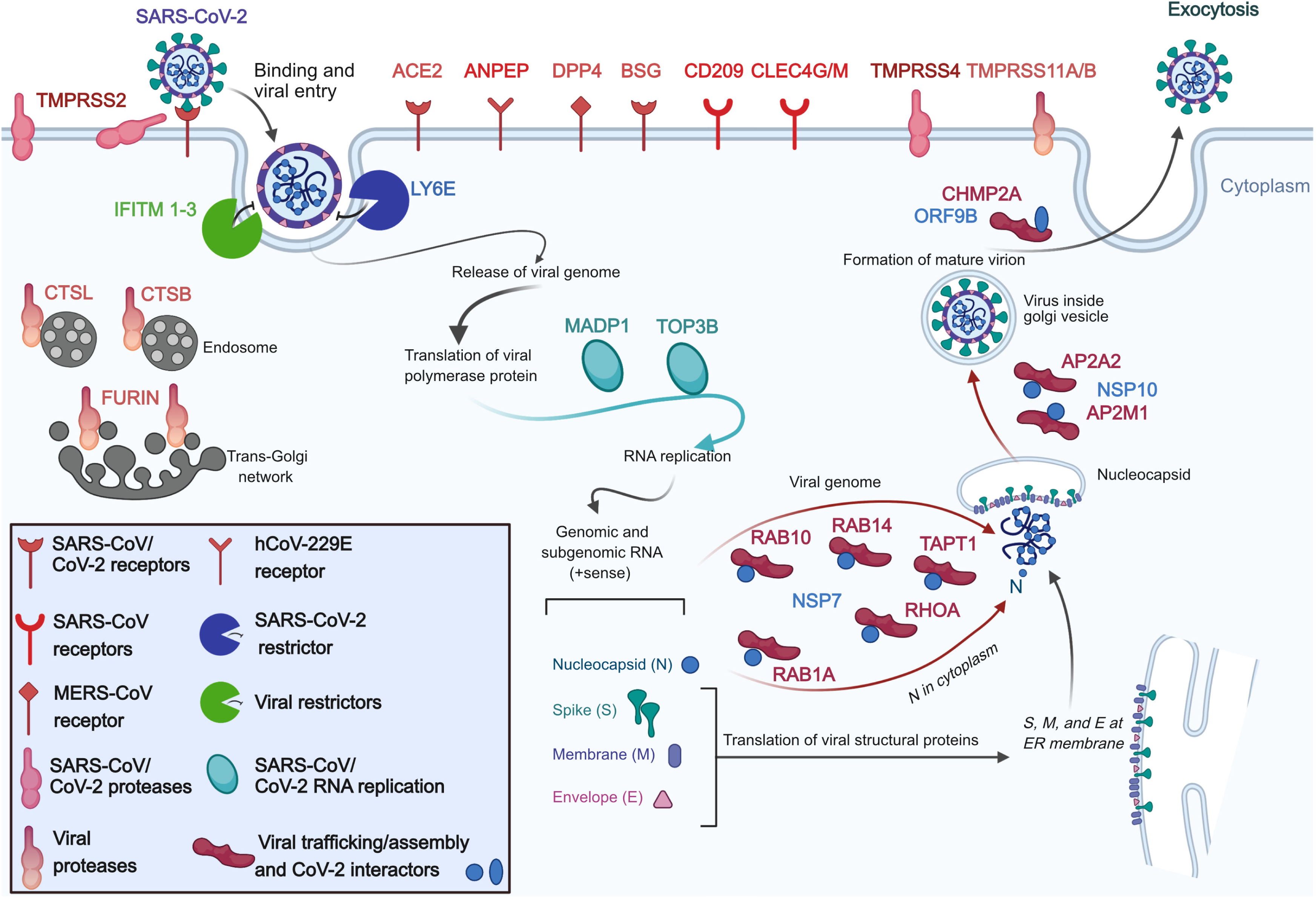
SARS-CoV-2 and coronavirus-associated receptors and factors (SCARFs) A cartoon illustration of the infection cycle of SARS-CoV-2 and its cellular interaction with entry factors (cell surface receptors, proteases, restriction factors) and post-entry factors (replication and assembly/trafficking factors) considered in this study. See text and Table S2 for further description and references.

## RESULTS

### SCARF curation

Because many cellular factors involved in viral replication, such as those involved in transcription, translation and other housekeeping functions, are unlikely to affect the tropism of the virus, we primarily focus on factors acting at the level of entry. Two factors have been established most rigorously to promote cellular entry of SARS-CoV-2 (and previously SARS-CoV) in human cells (Figure 1 and Table S2): the ACE2 receptor and TMPRSS2 protease (Hoffmann et al., 2020). BSG is a putative alternate receptor for both SARS-CoV and SARS-CoV-2, which has received experimental support (Chen et al., 2005; Wang et al., 2020b). We also included receptors which have been confirmed experimentally to facilitate entry of either SARS-CoV/hCoV-229E (ANPEP, CD209, CLEC4G/M) or MERS-CoV (DPP4), and are therefore candidates for promoting SARS-CoV-2 entry (Vankadari and Wilce, 2020). Next, we considered a number of cellular proteases, in addition to TMPRSS2, as alternative priming factors. TMPRSS4 was recently shown to be capable of performing this function for SARS-CoV-2 in human cells (Bertram et al., 2011; Zang et al., 2020). TMPRSS11A/B has been shown to activate the S peptide of other coronaviruses (Kam et al., 2009; Zmora et al., 2018). Additionally, FURIN is known to activate MERS-CoV and possibly SARS-CoV-2 (Mille and Whittaker, 2014; Walls et al., 2020) and Cathepsins (CTSL/B) can also substitute for TMPRSS2 to prime SARS-CoV (Simmons et al., 2013b). Importantly, we also enlisted several restriction factors that are known to protect cells against entry of SARS-CoV-2 (LY6E) (Pfaender et al., 2020) or a broad range of enveloped RNA viruses, including SARS-CoV (IFITM1-3) (Huang et al., 2011). We also considered a few additional factors that act post-entry, but relatively early in viral replication such as TOP3B and MADP1 (ZCRB1), which may expressed in a tissue/cell type-specific fashion and are known to be essential for genome replication of SARS-CoV-2 and SARS-CoV, respectively (Prasanth et al., 2020; Tan et al., 2012). Lastly, we included a set of proteins known to be involved in assembly and trafficking of a range of RNA viruses and have been shown recently to interact physically with SARS-CoV-2 structural proteins (Gordon et al., 2020), including members of the Rho-GTPase complex (RHOA, RAB10, RAB14, and RAB1A), AP2 complex (AP2A2 and AP2M1), and CHMP2A. In total, our list includes 28 SCARFs we deem solid candidates for modulating SARS-CoV-2 entry and replication in human cells (Figure 1).

### SCARF expression during pre-implantation embryonic development

To profile SCARF RNA expression in early embryonic development, we mined scRNA-seq data for human pre-implantation embryos (Yan et al., 2013). Our analysis revealed that *ACE2* mRNA is most abundant in the earlier stages of development, prior to zygotic genome activation (ZGA, 8-cell stage), indicating maternal RNA deposition (Figure 2A and S1A). *ACE2* transcript levels are depleted post-ZGA until the formation of the trophectoderm of the blastocyst in which they rise up again (Figure 2A and S1A). By contrast, *TMPRSS2* expression is only apparent in the primordial endoderm and trophectoderm lineages. In fact, none of the *TMPRSS* family members showed significant transcript levels (log2 FPKM >1) prior to ZGA (Figure 2A and Figure S1A-B). Pluripotent stem cells show high expression of *IFITM1-3* but no evidence of *ACE2* expression (Figure 2A and Figure S1A). Furthermore, analysis of another scRNA-seq dataset profiling ∼60,000 cells representing 10 cell types differentiated *in vitro* from pluripotent stem cells (Han et al., 2020) show no significant *ACE2* expression in any cells up to 20 days post-differentiation (Figure S1C). Together these data suggest that pluripotent stem cells and cells in early stage of differentiation are unlikely to be permissive to SARS-CoV-2 infection.

**Figure 2:**
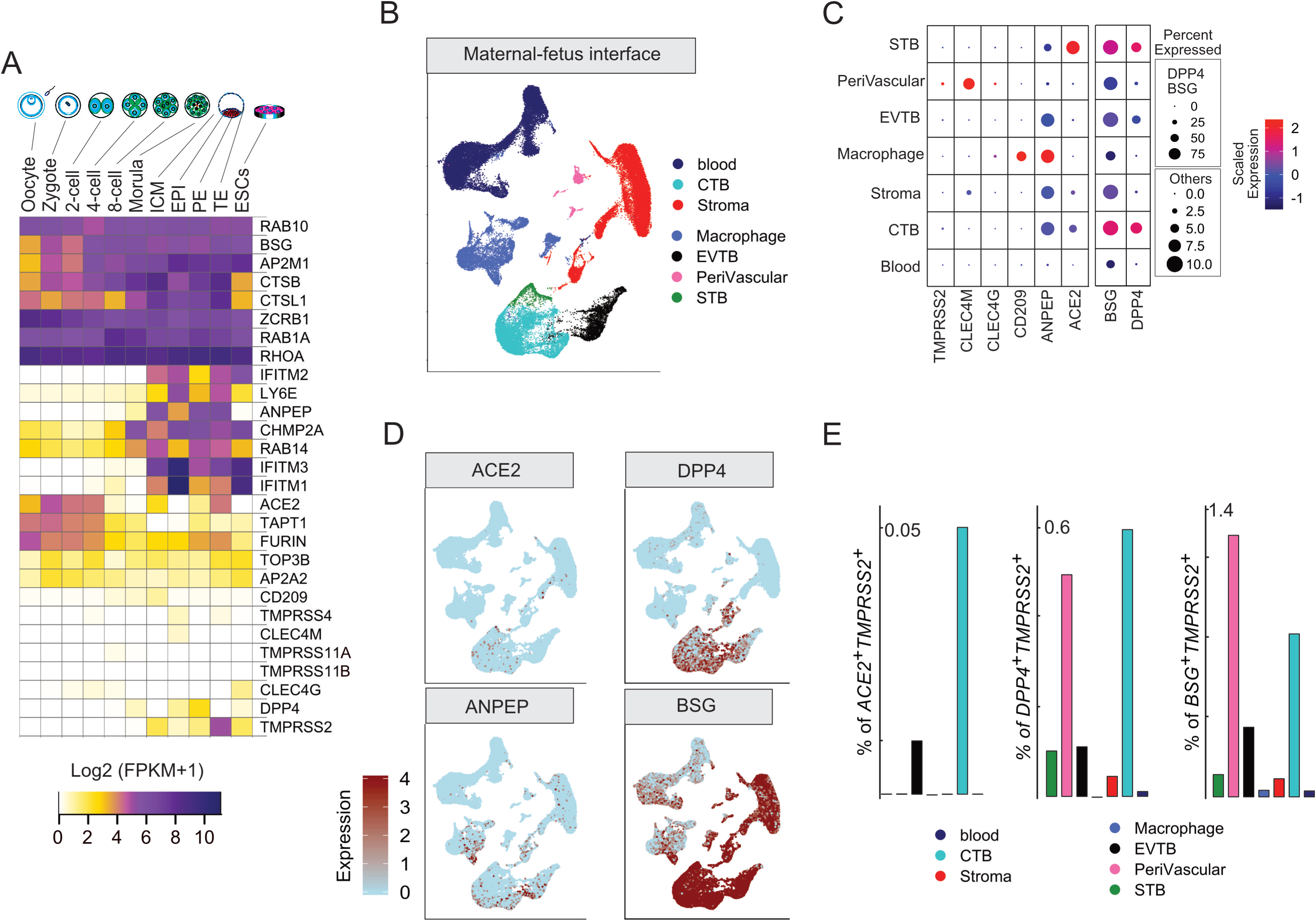
SCARF expression in preimplantation embryo and at the maternal-fetal interface. A. Heatmap of SCARF transcript levels for each stage of human preimplantation development. RPKM: reads per kilobase of transcript per million mapped reads. ICM: Inner Cell Mass, EPI: Epiblast, PE: Primitive Endoderm, TE: Trophectoderm, ESC: Embryonic Stem Cells. B. Uniform Manifold Approximation and Projection (UMAP) plot illustrating cell clusters identified at the maternal-fetal interface. Clusters consist of three trophoblast lineages (CTB: cytotrophoblasts, STB: syncytiotrophoblasts and EVTB: extravillous Trophoblasts), two decidual lineages (stroma and perivascular cells), immune cells (T cells, B cells, monocytes, dendritic cells, and natural killer cells), and macrophages (Hofbauer cells and M1 macrophages). See Figure S2 for a visualization of markers’ expression for each cell type. C. RNA transcript intensity and density of SCARFs (receptors and proteases) across the major cell types of the maternal-fetal interface. Dot colors correspond to the average Log2 expression level scaled to the number of unique molecular identification (UMI) values captured in single cells. The dot color scales from blue to red, corresponding to lower and higher expression, respectively. The size of the dot is directly proportional to the percent of cells expressing the gene in a given cell type. Since the percent of cells expressing the individual SCARF is much higher for DPP4 and BSG than for the other factors (“others”, i.e. ACE2, ANPEP, CD209, CLEC4G/M, and TMPRSS2), we use two different dot size scales for these two gene sets. D. Feature plots displaying unsupervised identification of cells expressing ACE2, DPP4, BSG, and ANPEP over the UMAP of maternal-fetal interface (see Figure 2B and S2). Cells are colored according to expression levels, from light blue (negligible to no expression) to red (high expression) (see Methods). E. Bar plots showing the percent of ACE2^+^TMPRSS2^+^ (left panel), DPP4^+^TMPRSS2^+^ (middle panel), and BSG^+^TMPRSS2^+^ (right panel) cells in the major cell types of the maternal-fetal interface (see also Table S3)

### SCARF expression at the maternal-fetal interface

The high level of *TMPRSS2*, *ACE2* and other coronavirus receptors such as *ANPEP* in the trophectoderm, which gives rise to the placenta, combined with low levels of *IFITMs* in this lineage (Figure 2A and Figure S1A) raises the possibility that the developing placenta may be vulnerable to SARS-CoV-2 infection. To investigate this, we turned to the transcriptomes of ∼70,000 single cells derived from tissues collected at the maternal-fetal interface during the first semester of pregnancy (Vento-Tormo et al., 2018), which include both embryo-derived cells (fetal placenta) as well as maternal blood and decidual cells. Our analysis of this dataset based on unsupervised clustering and examination of known markers recapitulated the major types of trophoblasts, decidua and immune cells (Figure 2B and S2A). Expression of *ACE2* and *DPP4* receptors was evident in cytotrophoblasts (CTB) and syncytiotrophoblasts (STB). *ANPEP* was abundantly expressed in all fetal lineages, while BSG was broadly expressed in maternal and fetal cells, but at much higher density in fetal cells (Figure 2C-D)*. CLEC4M* was the only potential receptor highly expressed in maternal-derived cells; our analysis identified this gene as a strong marker of decidual perivascular cells (Figure 2B-D). Interestingly, extravillous trophoblasts (EVT), which directly invade maternal tissue, showed low level of *ACE2* or *TMPRSS2*, but moderate to high levels of restriction factors *IFITM1-3* and *LY6E*, which were also expressed by immune and decidual cells (Figure S2B). *TMPRSS2*-expressing cells were comparatively less abundant within any cell types than those expressing receptors (Figure 2C). Thus, the maternal-placenta interface displays a complex pattern of SCARF expression.

To more finely assess the permissiveness of different placental cell types to SARS-CoV-2 entry, we quantified the fraction of each cell types co-expressing different combinations of receptors with proteases (predicted as more permissive) or with restriction factors (less permissive). The CTB stood out for having the largest fraction of cells double-positive for various receptor-protease combinations, including *ACE2^+^TMPRSS2^+^* (0.05%), *ACE2^+^FURIN^+^* (∼3%), *BSG^+^TMPRSS2^+^* (0.8%), *BSG^+^FURIN^+^* (10%), *DPP4^+^TMPRSS2^+^* (0.6%), and *DPP4^+^FURIN^+^* (∼10%) (Figure 2D-E, S2C and Table S4). Perivascular tissues also exhibited *BSG^+^TMPRSS2^+^* and *DPP4^+^TMPRSS2^+^* cells, albeit in fewer proportion compared to CTB (0.5 %) (Figure 2E and S2B). Interestingly, a substantial fraction of *DPP4*^+^ cells (∼20-80%) were co-expressing *IFITM1-3* and *LY6E* consistently across the whole dataset, whereas *ACE2*^+^ and *BSG*^+^ cells rarely co-expressed these restriction factors (Figure 2E, S2D and Table S4). Rather, *ACE2^+^* cells tend to co-express *TMPRSS2* and *FURIN,* but again this was mostly confined to a small subset of CTB cells (see also Figure 1D-E). Overall, these results suggest that the CTB is the cell type most susceptible to coronavirus infection within the first trimester placenta.

### SCARF expression in reproductive organs

Of all adult tissues surveyed via bulk RNA-seq by the GTEx consortium (Aguet et al., 2017), *ACE2* showed highest level of expression in human testis (Figure S4). To monitor more finely the expression profile of SCARFs in male and female reproductive tissues, we analyzed scRNA-seq datasets from testis samples collected from two healthy donors (Sohni et al., 2019) and adult ovary from five healthy donors (Wagner et al., 2020). In adult testis, we were able to recapitulate the expression clusters identified in the original report (Sohni et al., 2019), which consisted of early and late stages of spermatogonia (SPG), spermatogonial stem cells (SSCs), spermatids (ST), macrophages, endothelial and immune cells, each defined by a unique set of marker genes (Figure S3A-B). Turning to SCARFs, we observed that *TMPRSS2* is strongly expressed in both early and late SSCs, whereas CoV receptors are abundantly expressed in the early stage of SSCs (Figure 3A). Early SSCs expressing one of the receptors (*ACE2, BSG, DPP4, ANPEP*) were found to be consistently enriched for co-expression with *TMPRSS2* across the testis dataset (hypergeometric distribution, p-value <1e-7) (Figure 3B). Moreover, other SCARFs interacting with SARS-CoV-2 proteins and predicted to facilitate virus trafficking or assembly (Table S2) also show highest transcript levels in SSC and SPG (Figure 3C). In contrast, restriction factors were lowly expressed in all four clusters of spermatogonial cells (Figure 3A). Taken together these observations indicate spermatogonial cells may be highly permissive to SARS-CoV-2 infection.

**Figure 3:**
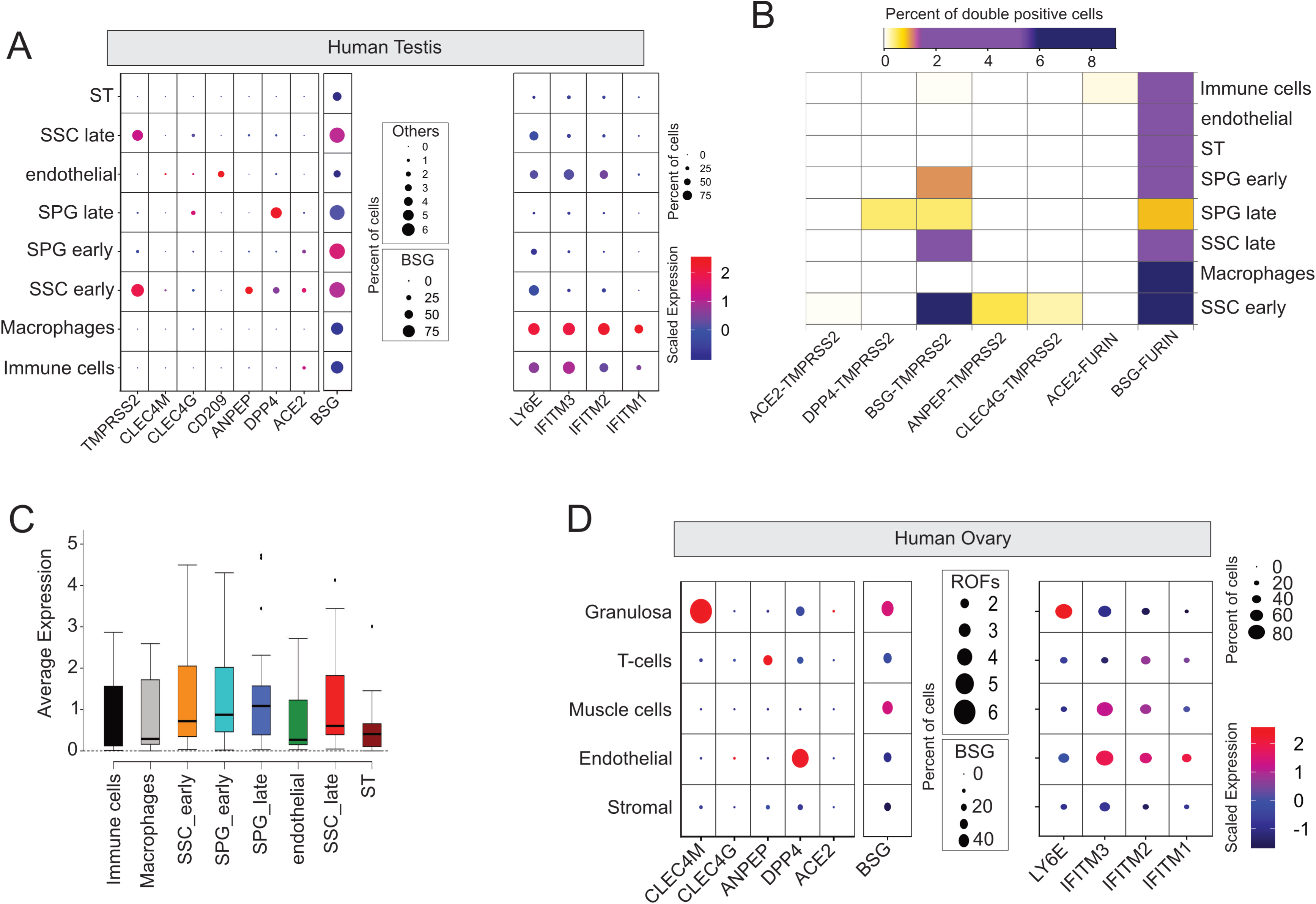
SCARF expression in reproductive organs. A. SCARF expression in the adult testis. See legend of Figure 2C for a description of dot plot display. Note that different dot size scales are used for BSG and “others” (i.e. ACE2, DPP4, ANPEP, CD209, CLEC4G/M, TMPRSS2). ST: spermatids, SPG: spermatogonia, SSC: spermatogonial stem cells. B. Heatmap of the fraction of double-positive cells for different receptor-protease combinations in each of the major cell types of the testis. Color scheme range from white for the lowest fraction (<0.05% of cells), gold for medium (0.05-2%), purple for high (2-6%), and dark blue for the highest fraction (>6%) of double-positive cells per cell type. C. Boxplots showing average single-cell expression of SCARFs in different testis cell types. D. SCARF expression in the adult ovary. Dot plot display as in Figure 3A. Note TMPRSS2 and TMPRSS4 transcripts were not detected (UMI = 0) in any single cell of the ovary.

Our analysis of ovarian cortex samples from 5 donors resolve the major cell types characteristic of this tissue, such as granulosa, immune, endothelial, perivascular, and stromal cells, each defined by a unique set of markers (Figure S3C-D), in concordance with the original article (Wagner et al., 2020). *ACE2*^+^ cells were generally rare across this dataset, and most evident in granulosa, where they show a relatively high level of expression per cell (Figure 3D). Alternate receptors were expressed at significant level in specific ovarian cell populations. For instance, *DPP4* was expressed in 4% of endothelial cells, while *CLEC4M* and *BSG* were highly expressed in granulosa (Figure 3D). T cells and endothelial layers were markedly enriched for *ANPEP* and *DPP4* transcripts, respectively (Figure 3D). Strikingly, however, we could not identify any single cell across the entire ovarian dataset with evidence of *TMPRSS2* expression, nor any of the alternate proteases *TMPRSS4, TMPRSS11A and TMPRSS11B*. This pattern was corroborated by examining the expression profile of all TMPRSS protease family members using bulk RNA-seq data from GTEx. None of these proteases appear to be expressed in ovarian tissue, whereas the testis expresses 7 out of 19 members including *TMPRSS2* (Figure S4 and S5). We note that the oocyte also lacks transcripts from any *TMPRSS* family members (see Figure 2A, S1A). Collectively, these data reveal a stark contrast between male and female reproductive organs: while early stages of spermatogenesis may be highly permissive for SARS-CoV-2 entry, the ovary and oocytes are unlikely to get infected.

### Human cell landscape reveals adult organs potentially most permissive to SARS-CoV-2

We analyzed a scRNA-seq dataset for ∼200,000 cells generated as part of the human cell landscape (HCL) project, which encompasses all major adult organs (Han et al., 2020). We favored this dataset over the Human Cell Atlas project (Rozenblatt-Rosen et al., 2017) because the HCL samples were prepared uniformly and processed through the same sequencing platform and raw counts for unique molecular identifiers (UMI) were publicly available, which enabled adequate normalization using our scRNA-seq analysis pipeline (see Methods). We selected 14 distinct adult tissues and clustered them into 33 distinct cell-types annotated using the markers described in the original article (Han et al., 2020) (Figure 4A and S6A).

**Figure 4:**
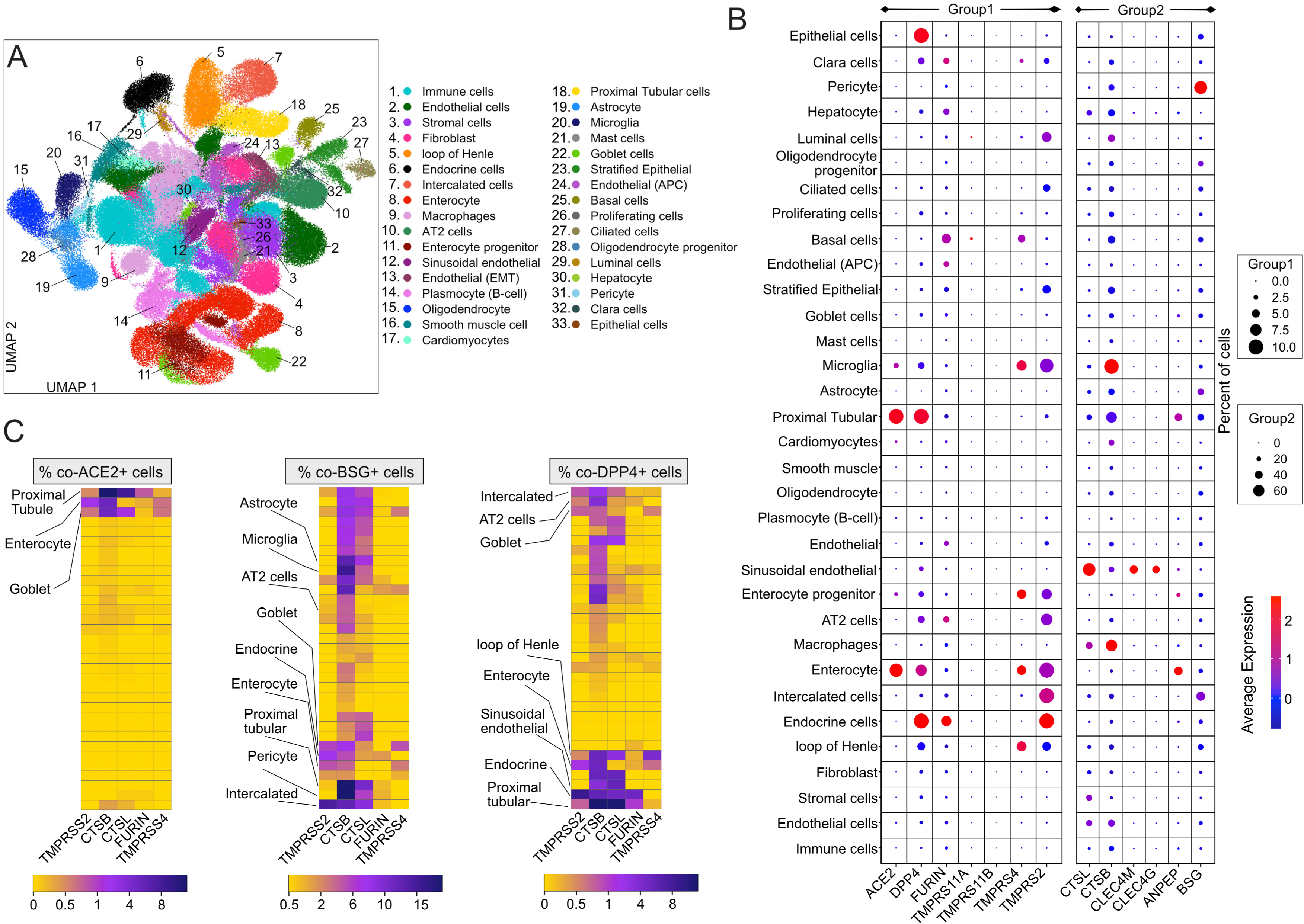
SCARF expression across 14 adult tissues. A. UMAP bi-plot resolving ∼200,000 single cells from 14 adult tissues obtained from the Human Cell Landscape into 33 different cell types. These cell types were consolidated from 78 cell clusters defined from the initial clustering (see also Methods and Figure S6 for tissues corresponding to each cell type). Markers for each cell type are listed in Table S4. B. Expression intensity and density of selected SCARFs (receptors and proteases) in each of the 33 cell types defined in Figure 1A. See legend of Figure 2C for a description of dot plot display. Different dot size scales are used for genes with low fraction of positive cells (group 1: up to 10%; ACE2, DPP4, FURIN, TMPRSS2, TMPRSS4, TMPRSS11A/B) and those with higher fraction (group 2: up to 60%; BSG, ANPEP, CLEC4G/M, CTSB/L). C. Heatmap of the fraction of double-positive cells for different receptor-protease combinations in each of the 33 cell types.

We first focus our analysis on *ACE2* and *TMPRSS2*, the two genes most robustly established as entry factors for SARS-CoV-2 (Table S2). Transcripts for both genes were detected (> 0.1 % of total cells) primarily in colon, intestine (ileum, duodenum and jejunum), gallbladder, and kidney cells (Figure S6B-D). We detected little to no expression of *ACE2* and/or *TMPRSS2* in the remaining tissues and cell types (Figure 4B), including type 1 (AT1) and type 2 (AT2) alveolar cells of the lung. This latter observation is at odds with earlier reports (Table S1), but in line with a recent study using a wide array of techniques to monitor *ACE2* expression within the lung, including transcriptomics, proteomics and immunostaining (Hikmet et al., 2020). Cardiomyocytes (from the heart) show relatively high levels of ACE2, but lacked TMPRSS2 expression (Figure 4B and S6B-E). It is also important to emphasize that even when *ACE2* and/or *TMPRSS2* were detected at appreciable RNA levels in a given organ or cell type, only a very small fraction of cells was found to express both genes simultaneously. For instance, the kidney shows relatively high levels of *ACE2* and *TMPRSS2*, but only 0.01% cells were double-positive for these factors. Only three cell types stood out in our analysis for relatively elevated levels of co-expression of *ACE2* and *TMPRSS2* (0.5-5% of cells double-positives): enterocytes, proximal tubule cells, and goblet cells (Figure 4B-C), which we will return to below.

Examining alternative receptors revealed a more complex picture, in part reflecting the tissue-specificity of these genes. For instance, *CLEC4G* and *CLEC4M* were highly expressed in the liver as previously reported (Domínguez-Soto et al., 2009), and more specifically in sinusoidal endothelial cells and hepatocytes (Figure 4B and S6B-C). *BSG* marked the pericytes and astrocytes of the cerebellum, as well as intercalated cells of the kidney, whereas *CD209* was enriched in macrophages (Figure 4B and S6B). *ANPEP* and *DPP4* were often co-expressed in multiple organs and cell types, including prostate, lung, kidney, colon and small intestine (Figure S6B). Furthermore, many of the same organs exhibited a significant fraction (∼2 to ∼8%) of cells triple-positive for *ANPEP*, *DPP4* and *TMPRSS2*, albeit in variable amounts (Figure S6A-C). The prostate showed particularly high expression level of *DPP4* (∼15% double-positive with *TMPRSS2* or *FURIN*) and moderate levels of *ANPEP* (∼5% double-positive with *TMPRSS2* or *FURIN*) (Table S5). Expression of these factors within the prostate was primarily driven by endocrine cells, which displayed the highest fraction of *ANPEP^+^DPP4^+^TMPRSS2^+^* cells among all the cell types defined in our analysis (Figure 4B-C, S6A-C and Table S5). Within the lung, AT2 cells also prominently expressed *DPP4, BSG* and *ANPEP*, along with *TMPRSS2* and/or *FURIN* (Figure 4B-C), suggesting that these receptors, rather than *ACE2*, could represent the initial gateway by which SARS-CoV-2 infect AT2 cells, which are known to be extensively damaged in SARS and COVID-19 pathologies (Qi et al., 2020). Intercalated cells of the kidney as well as goblet cells and enterocytes of the colon were also highly enriched for *TMPRSS2^+^DPP4^+^BSG^+^* cells (Figure 4B-C and S6A-C).

The small intestine was rather unique among the organs represented in this dataset for expressing high levels of *ACE2, ANPEP* and *DPP4* (Figure S5B and S6B-C), with highest levels in the jejunum, which also exhibited copious amount of *TMPRSS2* (Figure S6 C-E and S7). This is in slight disagreement with another study, which analyzed the Human Cell Atlas data and suggested that the ileum displayed the highest level of *ACE2* transcripts (Ziegler et al., 2020). Consistent with this study, however, we found that the bulk of expression of these factors in the small intestine is driven by enterocytes and their progenitors (Figure 4A-C and S5, S6B and S7). Only two other cell types were equally remarkable for expressing the quadruple combination of *ACE2, ANPEP, DPP4*, and *TMPRSS2*: (i) goblet cells, epithelial cells lining and producing mucus for several organs including duodenum, ileum, colon and gallbladder, and (ii) proximal tubular cells of the kidneys (Figure 4A-C and S6A-B, S7A-D). In addition, the enterocytes, and goblet cells were enriched for SCARFs interacting with SARS-CoV-2 proteins and implicated in virus trafficking or assembly (Figure S7E).

In summary, coronavirus entry factors appear to be expressed in a wide range of adult healthy organs, but in restricted cell types, including pericytes, astrocytes, and microglia of the cerebellum, sinusoidal endothelium of the liver, endocrine cells of the prostate, enterocytes of the small intestine, goblet cells, and the proximal tubule of the kidney.

### Nasal epithelium

The nasal epithelium is thought to represent a major doorway to SARS-CoV-2 infection (Sungnak et al., 2020; WU et al., 2020). Since this tissue was not included in the HCL dataset, we analyzed another scRNA-seq dataset derived from six nasal brushing samples from healthy donors collected by three independent studies (Deprez et al., 2019; Garcıá et al., 2019; Vieira Braga et al., 2019) (Table S3). Our analysis of this merged dataset reveals four major cell clusters consistent across all 6 samples, corresponding to ciliated, secretory, suprabasal epithelial cells and natural killer cells (Figure 5A, Figure S8). Natural killer cells express mostly *DPP4*, while the three epithelial cell types showed low to moderate RNA amounts of *ACE2*, *ANPEP, BSG* and *TMPRSS2* (Figure 5A, Table S6). *ACE2* RNA was most abundant in ciliated cells, while *ANPEP* was most highly expressed in secretory cells. Conversely, *BSG* was abundant throughout all the nasal epithelial cell types, albeit at higher level in suprabasal cells (Figure 5A, S9A and Table S6). *TMPRSS2* was also expressed in all three nasal epithelial cell types, with highest density in ciliated cells (41%). Overall, the percentage of *ACE2^+^TMPRSS2^+^* cells was relatively low within each cell type (2.3% in ciliated, 1.6% in secretory and 1% in suprabasal cells) (Table S6). In contrast with the digestive system (characterized above, Figure 4B-C), nasal epithelial cells rarely show co-expression of *ANPEP, DPP4* and *ACE2*, but rather exclusive expression of one of these three receptors (Figure 5A and S9A). By contrast, restriction factors *IFITM3* and *LY6E* were robustly expressed in all three nasal epithelial cell types (Figure 5A) with highest levels in secretory and suprabasal cells (Table S6). Lastly, we calculated the percentage of double/triple positive cells for various combinations of *ACE2*, *TMPRSS2* and restriction factors (Table S6). We found that 85% and 65% of *ACE2^+^TMPRSS2^+^* ciliated cells are also positive for *LY6E* and *IFITM3*, respectively (Table S6). Together these data suggest that the nasal epithelium expresses various combinations of factors that in principle could facilitate SARS-CoV-2 infection, but it also expresses robust basal level of restriction factors, which may act as a strong protective barrier in this tissue.

**Figure 5:**
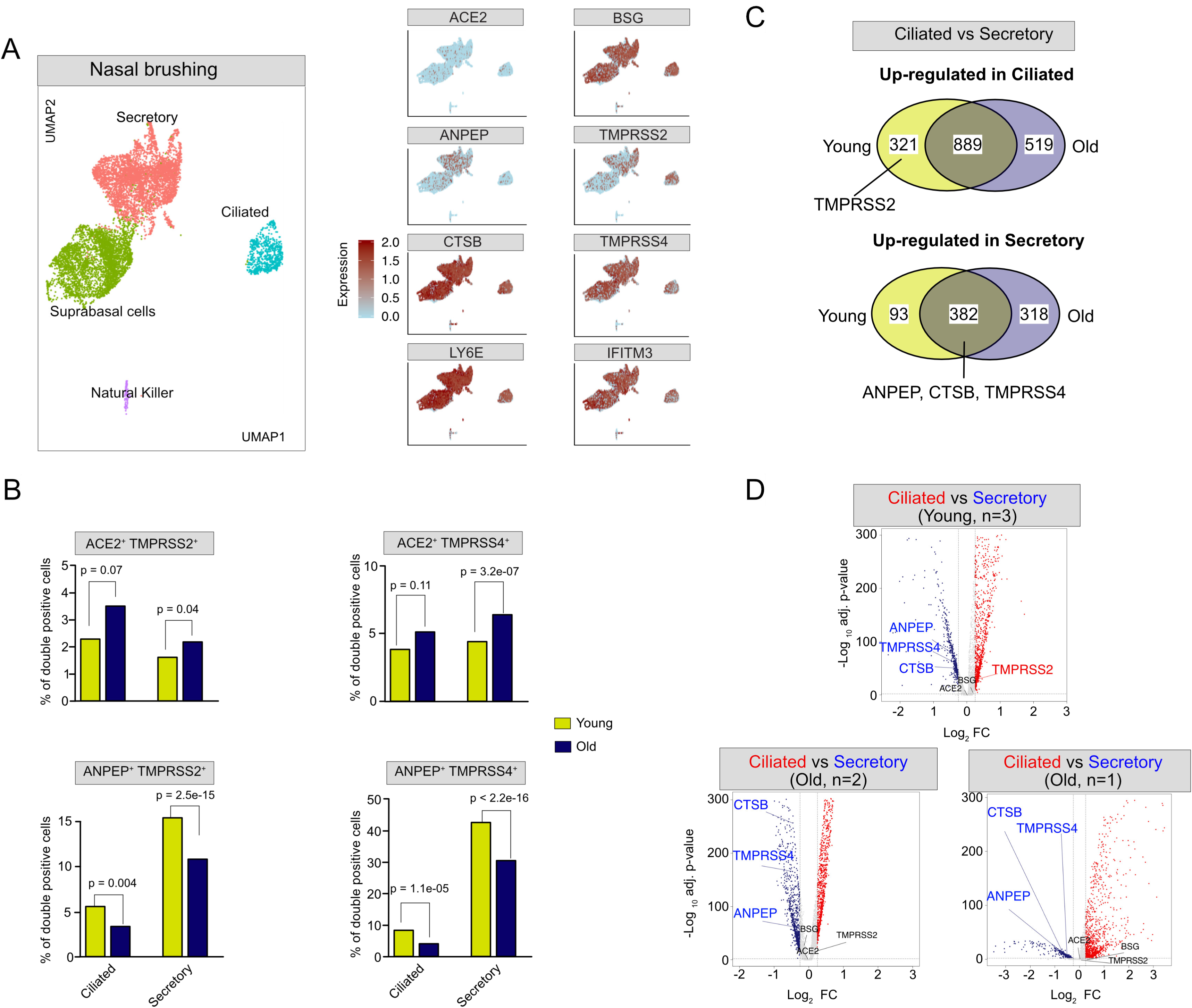
SCARF expression in nasal brushing samples. A. Left panel: UMAP bi-plot shows the cell clustering of 6 nasal epithelial scRNA-seq samples from 3 independent studies (See also Figure S8). Right panel: Feature plots showing expression of hCoV receptors (*ACE2, ANPEP*, and *BSG*), proteases (*TMPRSS2, CTSB*, and *TMPRSS4*), and restriction factors (*LY6E* and *IFITM3*) over the UMAP plot. Cells are colored according to expression levels, from light blue (negligible to no expression) to red (high expression) (see Methods). B. Bar plots comparing the percent of double-positive cells for different receptor/protease combinations in ciliated and secretory cells of the young (N=3) and old (N=3) nasal samples. A one-sided Fisher’s exact test was employed to test the significance of the differences observed between the young and old groups. C. Venn diagrams illustrating shared and unique differentially expressed genes (DEGs, adjusted-*p*-value <0.01, and average log fold change |0.25|) between ciliated and secretory cells within young (N=3) and old (N=3) samples. The upper panel shows genes up-regulated in ciliated cells, while the lower panel shows genes up-regulated in secretory cells. D. Volcano plots showing DEGs between ciliated and secretory cells identified independently for each of the three studies. All detected genes are plotted in a two-dimensional graph with x-axis representing the log-fold change in transcript levels (calculated with the ‘Seurat’ package, see Methods) and the y-axis representing the significance level (-log10 adjusted p-value, Wilcoxon Rank Sum test, adjusted with Bonferroni correction). Higher value on the y-axis denotes stronger significance levels.

### Age may modulate SCARF expression in the nasal epithelium

To investigate a possible age-effect in the expression of CoV entry factors within the nasal epithelium, we took advantage of the fact that three of the samples analyzed above were collected from relatively young donors (24-30 years old), while the other three came from older individuals (50-59 years old) (Figure S8 and Table S3). While this is a small sample, this enabled us to split the data into a ‘young’ and ‘old’ group and compare the percentage of secretory and ciliated cells positive for entry factors between the two groups (Table S3). The percentage of cells positive for either *ACE2, TMPRSS2* or *TMPRSS4* was comparable between the two groups and these factors were most highly expressed in ciliated cells of both groups. However, the percentage of double-positive cells (*ACE2^+^TMPRSS2^+^* or *ACE2^+^TMPRSS4*^+^) were significantly higher in the old group, both within ciliated and secretory cell populations (Figure 5B, S9B and Table S6). Interestingly, the percentage of *ANPEP^+^TMPRSS2^+^* or *ANPEP^+^TMPRSS2^+^* double positives cells showed the opposite trend: they were significantly more frequent in the young group, both within ciliated and secretory cells (Figure 5B). To examine whether these differences were driven by an age-dependent shift in the relative expression of receptors and/or proteases during cell differentiation within the nasal epithelium, we conducted a global differential gene expression analysis between ciliated and secretory cells within each age group and each independent study. We found that *ANPEP*, *TMPRSS4* and *CTSB* were significantly up-regulated in secretory cells relative to ciliated cells in all three studies, regardless of age (Figure 5C). Conversely, *TMPRSS2* levels were up-regulated in ciliated cells of the young group, but remains unaltered in old individuals regardless of the study of origin (Figure 5C-D, Table S7). These results suggest that there is a shift in *TMPRSS2* regulation during nasal epithelium differentiation in young individuals that is not occurring in old individuals (Figure 5C and 5D, Table S7).

### Conservation of SCARF expression across primates

To examine the evolutionary conservation of SCARF expression across primates, we analyzed a recently published comparative transcriptome dataset (bulk RNA-seq) of heart, kidney, liver, and lung primary tissues from human (N=4), chimpanzee (N=4), and Rhesus macaque (N=4) individuals, comprising a total of 47 samples (Blake et al., 2020). As expected, UMAP analysis showed that the samples clustered first by tissue, then by species (Figure 6A).

**Figure 6:**
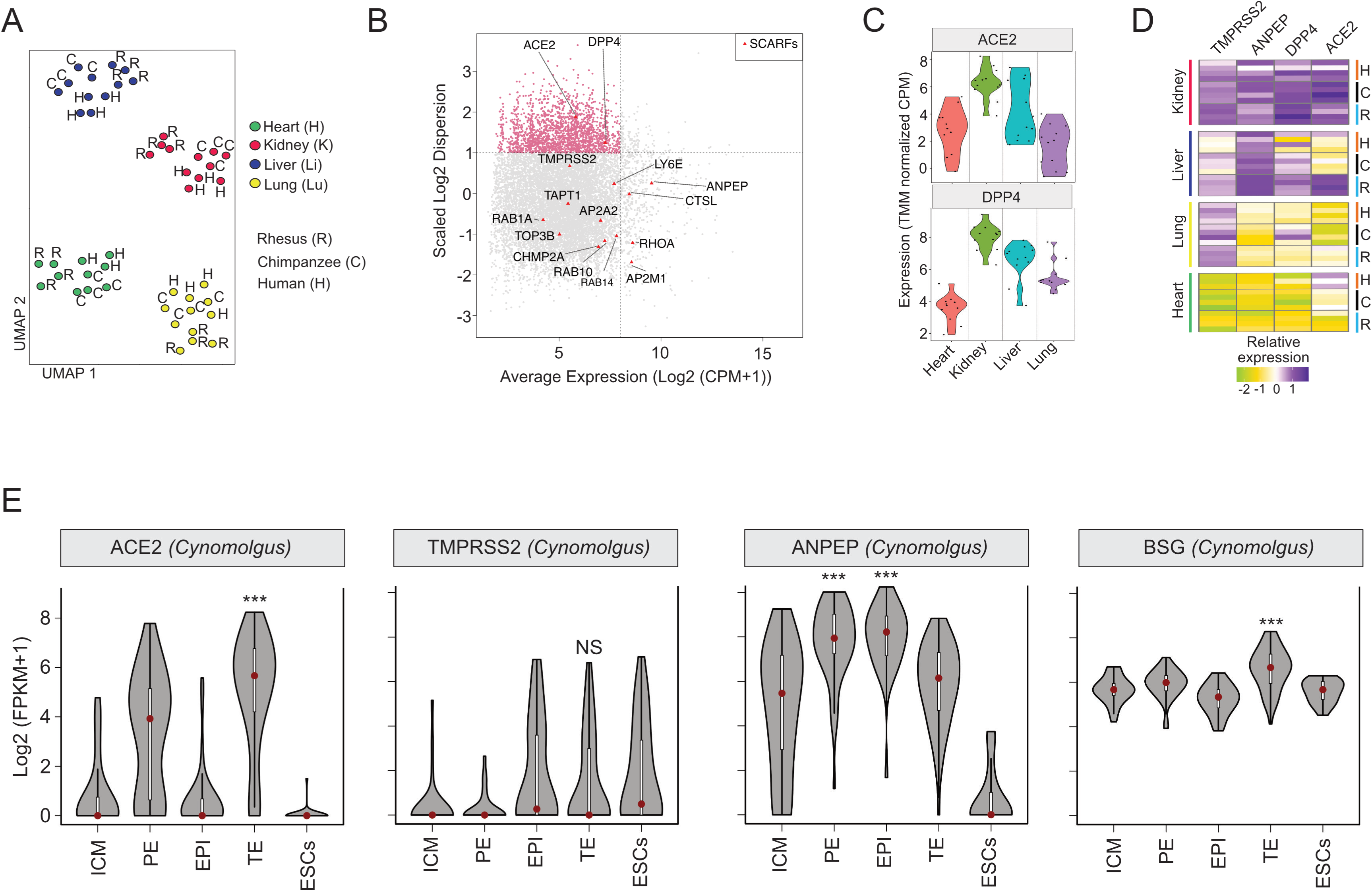
Cross-species analysis of SCARF expression. A. UMAP clustering of 47 individual RNA-seq samples derived from four tissues (heart, lung, liver, and kidney) of three species (human, chimpanzee, Rhesus macaque) based on 11,929 orthologous RefSeq genes most variable in expression across the samples (see Methods). B. Scatterplot of normalized mean expression in CPM (Counts Per Million) of each orthologous gene (x-axis) and scaled dispersion (y-axis) across the 47 samples. Every point corresponds to a single gene. The most 1,987 variable genes across the samples (Log2 scaled dispersion >1) are marked as pink dots. Red triangles mark SCARFs. C. Violin plot showing expression dispersion of ACE2, TMPRSS2, DPP4, and ANPEP levels per tissue (TMM-normalized CPM). Each dot comes from a different sample. D. Heatmap of scaled expression levels (TMM-normalized CPM) of ACE2, TMPRSS2, ANPEP, and DPP4 for each of the individual samples (N=47) grouped by tissues and species. E. Violin plots showing Log2-normalized expression profile of ACE2, TMPRSS2, and ANPEP in the preimplantation blastocyst of Cynomolgus macaque obtained by scRNA-seq.

Overall, we observe a low level of tissue-specificity and high level of inter-specific conservation in the expression of most SCARFs, with the notable exceptions of *ACE2* and *DPP4* receptors, which were included amongst the most variably expressed genes across tissues and species (Figure 6B). Specifically, liver and lung samples from chimpanzee showed relatively high *DPP4* expression but very low *ACE2* expression relative to human and macaque (Figure 6C-D). Liver and lung samples from human and chimpanzee did not express *ACE2*, however the macaque liver showed high expression level for *ACE2*, *DPP4, ANPEP, and TMPRSS2* genes (Figure 6D). These results suggest that macaques may be more prone to liver coronavirus infection than humans or chimpanzees. In agreement with our scRNA-seq analysis of the HCL dataset, we found that *ACE2, DPP4, ANPEP* and *TMPRSS2* genes are all expressed at higher level in the kidney of all three primates, relative to the other three organs. Thus, out of the four organs examined in this analysis, the kidney appears to be the most readily permissive to coronavirus infection in all three species (Figure 6C-D).

Finally, we analyzed the pattern of SCARF expression in the blastocyst of Cynomolgus macaques (a close relative of the Rhesus macaque) in comparison to that of the human blastocyst (Figure 2A and S1A). Intriguingly, the two species show distinct expression patterns for several entry factors. *TMPRRS2* was highly expressed in the human trophectoderm (TE), but transcripts for this gene were essentially undetectable in the macaque blastocyst, including TE (Figure 6E). Also, *ANPEP* was downregulated in the macaque TE compared with the rest of the blastocyst lineages, while it was upregulated in the human TE (compare Figure S1A and Figure 6E). Thus, there may be substantial differences in the susceptibility of human and macaque early embryos to coronavirus infection.

## DISCUSSION

COVID-19 has created a formidable health and socioeconomic challenge worldwide and an urgency to develop treatments and vaccines. A prerequisite to the success of these endeavors is to gain a better understanding of the biology and pathology of SARS-CoV-2, a newly emerged coronavirus responsible for millions of infections and over 300,000 deaths as of mid-May 2020. While it is clear that COVID-19 is primarily a respiratory disease which causes death via pneumonia, many unknowns remain as to the extent of tissues and cell types vulnerable to SARS-CoV-2. How host genetic factors interact with the virus and modulate the course of an infection also remain poorly understood. Our study, along with several others (published or preprinted over the past few weeks, Table S1) have tapped into vast amount of publicly available scRNA-seq data to profile the expression of host factors thought to be important for entry of SARS-CoV-2 in healthy tissues. Because the basal expression level of these factors determines, at least in part, the tropism of the virus, this information is foundational to predict which tissues are more vulnerable to infection. These data are also important to guide and prioritize clinical interventions and pathological studies, including biopsies. Finally, this type of analysis has the potential to reveal possible routes of infection within and between individuals.

Our study distinguishes itself from all other studies reported thus far (see Table S1) by the wider range of factors (SCARFs) examined across a large array of tissues. Many studies focused exclusively on the respiratory tract (Sungnak et al., 2020; Zhao et al., 2020) or other individual organ systems such as the brain, the olfactory system, or the GI tract (see Table S1). We interrogated a wide and unbiased set of organs, largely conditioned by public availability of raw data (e.g. UMI counts) as to apply uniform normalization procedures across datasets. For instance, we were able to integrate HCL samples that were prepared and processed through a single sequencing facility and platform. Also, it is important to emphasize that only two datasets examined here (placenta and nasal cavity from the Human Cell Atlas) were previously analyzed elsewhere (Sungnak et al., 2020). Thus, our analyses provide an independent replication of findings reported elsewhere for some tissues (see Table S1) as well as a trove of new observations (detailed below). The ultimate goal is to deliver within a single study a broad and robust foundation for future investigations, as well as a user-friendly searchable interface.

A unique asset of this study lies in the range of SCARFs integrated in the analysis (Table S2). Most other studies have focused exclusively on *ACE2* and/or *TMPRSS2* (see Table S1). A handful of studies considered one or a few additional receptors (e.g. *DPP4*, *ANPEP*) in a subset of their analyses (Sungnak et al., 2020). To our knowledge, no other study has included all the human coronavirus receptors considered here, even though these alternate receptors have been experimentally confirmed to interact with and facilitate entry of SARS-CoV-2 (*BSG*) or the closely related SARS-CoV (*CD209*, *CLEC4G/M*) (see Table S2 for references). Furthermore, our study is the first to consider factors known or predicted to restrict SARS-CoV-2 at the level of entry, such as LY6E and IFITMs. While these restriction factors are known to be interferon-inducible (Jia et al., 2012; Mar et al., 2018), their basal level of expression is likely to be a key determinant of SARS-CoV-2 tropism. Indeed, it is well established that coronaviruses are equipped with multiple mechanisms to suppress interferon and other immune signaling pathway (Frieman and Baric, 2008; Gralinski and Baric, 2015) and there is growing evidence that SARS-CoV-2 infection also illicit a ‘muted’ interferon response (Blanco-Melo et al., 2020; Wyler et al., 2020). As such, the basal level at which restriction factors are expressed in a given target cell may be the major obstacle the virus encounters at the onset of an infection.

Below we summarize and contrast our findings to existing studies published or available in preprint repositories as of May 2020 (listed in Table S1) and discuss how our results further our understanding of SARS-CoV-2 tropism and pathology.

### The developing embryo may be at risk of infection

To our knowledge, there has been no prior description of SCARF expression during pre-implantation development. While we detect high amount of *ACE2* mRNA in the zygote and up to the 4-cell stage, it precipitously drops at ZGA and remained at very low or undetectable levels in the pluripotent cells of the early embryo. *ACE2* remains silent in pluripotent stem cells in culture, even after 20 days of in-vitro differentiation to various cell types. These observations suggest that *ACE2* RNA is heavily deposited in the zygote (presumably from the oocyte), but the gene is subsequently silenced in stem cells and poorly differentiated cells. The idea that *ACE2* expression positively correlates with the level of cellular differentiation has been observed in the context of the airway epithelium (Jia et al., 2005).

The only pre-implantation lineage with substantial *ACE2* expression is the trophectoderm, where *TMPRSS2* is also highly and consistently expressed. During embryonic development, the trophectoderm gives rise to all the different types of trophoblasts (placental fetal cells). Accordingly, we observe that *ACE2* and *TMPRSS2* expression persists in a subset of trophoblasts at least up to the first trimester of pregnancy. Our results extend recent observations based on an independent analysis of the same placenta dataset, which remarked *ACE2* expression in certain cell types of placenta/decidua (Sungnak et al., 2020). We further these observations by identifying a small population of cytotrophoblasts co-expressing *TMPRSS2* with *ACE2, BSG* and/or *DPP4,* but exhibiting very low levels of *IFITM* and *LY6E* restriction factors. We conclude that a subset of trophoblast cells may be permissive to SARS-CoV-2 entry. Overall, our findings for the placenta are consistent with current clinical data suggesting that vertical transmission of SARS-CoV-2 from infected mother to fetus is plausible, but probably rare (Baud et al., 2020; Chen et al., 2020a; Cui et al., 2020; Hosier et al., 2020; Zeng et al., 2020). Future studies should be directed at examining SCARF expression at different stages of pregnancy and evaluating whether SARS-CoV-2 infection could compromise pregnancy.

### Ovarian cells may be resistant to SARS-CoV-2, but spermatogonial cells seem highly permissive

While we found no evidence for expression of any TMPRSS genes considered in female reproductive tissues, our analysis suggests that the early stages of spermatogenesis are vulnerable to SARS-CoV-2 and likely other coronaviruses. Indeed, we observe that spermatogonial cells (and stem cells) express high level of *ACE2, TMPRSS2, DPP4, ANPEP*, and low level of *IFITM* and *LY6E* restriction factors. To our knowledge, these observations have not been reported elsewhere. One study reported high level of *ACE2* expression in spermatogonial, Sertoli and Leydig cells (Wang and Xu, 2020), but did not investigate other SCARFs. Another study reported low level of *ACE2* and *TMPRSS2* in spermatogonial stem cells (Sungnak et al., 2020), but analyzed a different dataset apparently depleted for spermatogonial cells (data not shown). While it is unknown whether SARS-CoV-2 or other coronaviruses can infiltrate testes, it is notable that post-mortem autopsy of male patients infected by SARS-CoV revealed widespread germ cell destruction, few or no spermatozoon in seminiferous tubules, and other testicular abnormalities (Xu et al., 2006). These and other observations (Fanelli et al., 2020; Wang et al., 2020c), together with our finding that prostate endocrine cells also appear permissive for SARS-CoV-2, call for pathological examination of testes as well as investigation of reproductive functions in male COVID-19 patients.

### Respiratory tract: how does SARS-CoV-2 infect lung cells?

Because COVID-19, as well as SARS, are primarily respiratory diseases, the lung and airway systems have been extensively profiled for *ACE2* and *TMPRSS2* expression (Table S2), two SCARFs believed to be primary determinant for SARS-CoV-2 tropism. Paradoxically, healthy lung tissues as a whole show only modest expression for either *ACE2* and *TMPRSS2*, which is readily apparent in a number of expression databases and widely available resources such as GTEx (Figure S4). Nonetheless, a number of studies mining scRNA-seq data reported marked expression of *ACE2* and/or *TMPRSS2* in a specific lung cell population called alveolar type II (AT2) cells (Chow and Chen, 2020; Travaglini et al., 2019; Zhao et al., 2020). However, these observations have been challenged by more recent studies. For instance, Hikmet et al. could not confirm ACE2 expression in AT2 cells, neither through re-analysis of 3 different lung scRNA-seq datasets, nor by immunohistochemistry (Hikmet et al., 2020). Ziegler et al. analyzed a different lung scRNA-seq dataset and found only a very small fraction (>1%) of AT2 cells expressing both *ACE2* and *TMPRSS2* transcripts (Ziegler et al., 2020). Likewise, Aguiar et al. detected only very low amount ACE2 RNA or protein in alveoli and airway epithelium, but showed that the alternative receptor BSG was consistently expressed in respiratory mucosa from 98 human samples (non-COVID-19) (Aguiar et al., 2020). Our results, which stem from an independent analysis of yet another scRNA-seq dataset (Han et al., 2020) are consistent with the notion that basal *ACE2* RNA levels in lung cells, including AT2 cells, are very low. Importantly, we observed that AT2 cells do co-express the putative alternate receptors *BSG*, *ANPEP* and/or *DPP4* along with *TMPRSS2* at appreciable frequency (0.2-0.7%). Taken together, these data question whether ACE2 is the primary receptor by which SARS-CoV-2 initiates lung infection. A non-mutually exclusive explanation is that *ACE2* expression is widely variable in the lung due to genetic or environmental factors. Consistent with the latter, evidence is mounting that *ACE2* can be induced by interferon and other innate immune signaling (Ziegler et al., 2020).

### CNS and Heart: do the clinical symptoms manifest CoV-2 infection?

Some COVID-19 patients show neurological symptoms (De Felice et al., 2020) such as encephalitis, strokes, seizures, loss of smell, but it remains unclear whether SARS-CoV-2 can actively infect the CNS. In one case diagnosed with encephalitis, SARS-CoV-2 RNA was detected in the cerebellar spinal fluid (Moriguchi et al., 2020). Some have reported *ACE2* expression in various brain cells (Chen et al., 2020b). We observed *ACE2* and *TMPRSS2/4* expression specifically in microglial cells, although the two genes were rarely co-expressed within the same cells. We also found that the alternate receptor *BSG* was abundantly expressed in pericytes and astrocytes, but in those cell types *TMPRSS2/4* was apparently not expressed. However, *FURIN* and *CTSB* proteases were often co-expressed with *BSG* in these cells, suggesting potential alternate routes of viral entry in those cell types. Because pericytes are located in the vicinity of the blood-brain barrier, these cells may act as a gateway to CNS infection.

Whether the heart can be infected by SARS-CoV-2 is another open question. Severe heart damage and abnormal blood clotting has been reported in a substantial fraction of COVID-19 patients (Shi et al., 2020; Wang et al., 2020a). We and others (Litviňuková et al., 2020) found that *ACE2* is expressed in cardiomyocytes, but the same cell population does not appear to express *TMPRSS2/4*, so it remains unclear how the virus could infiltrate cardiomyocytes. Nonetheless, we observe that *FURIN* was co-expressed with *ACE2* in a very small fraction (<0.1%) of cardiomyocytes. While it is unclear whether SARS-CoV-2 can use FURIN to prime infection (Litviňuková et al., 2020), our findings suggest a possible path to heart infection.

### Nasal epithelium: niche or battleground?

Recently, Sungnak et. al. showed that SARS-CoV-2 entry factors are highly expressed in secretory and ciliated cells of the nasal epithelium (Sungnak et al., 2020). In agreement, we found that the percentage of *ACE2*^+^ or *TMPRSS2^+^* cells was higher among ciliated cells than among secretory or suprabasal cells. Conversely, we found that the percentage of *ANPEP^+^* cells was higher among secretory or suprabasal cells than in ciliated cells, while *BSG* was broadly expressed throughout the nasal epithelium. We also note that 19% of suprabasal cells expressed *TMPRSS2*. Yet, the percentage of *ACE2^+^TMPRSS2^+^* cells remained rather low across the nasal epithelium while *IFITM3* and *LY6E* restriction factors showed high RNA levels throughout this tissue. This latter finding is consistent with the observation of Sungnak et al. (2020) that *ACE2* expression in goblet cells correlates with that of immune genes and antiviral factors, though their study did not single out these specific restriction factors. Collectively these observations point to the nasal epithelium as an early battleground for SARS-CoV-2 infection, the outcome of which may be critical for the pathological development of COVID-19.

It is clear that COVID-19 causes more severe complications in patients with advanced chronological age. To our knowledge, the expression dynamics of SCARFs in relationship with age has not been explored in depth yet. One study reported that, paradoxically, *TMPRSS2* expression levels tend to mildly decrease with age in human lung tissue (Chow and Chen, 2020). We found that neither *ACE2* nor *TMPRSS2/4* on their own were differentially expressed between young and old nasal epithelia, but we observed that the percentage of *ACE2^+^TMPRSS2^+^/4^+^* double-positive cells was significantly greater in older donors, both within ciliated and secretory cells. Conversely, the percentage of *ANPEP*^+^*TMPRSS2^+^/4^+^* cells was significantly higher in younger donors. Considering ANPEP association with SARS-CoV infection (Kong et al., 2009; Yu et al., 2003), these trends fit with the observations that the median age for COVID-19 patients in China has been reported at 51 years old (Aylward, Bruce (WHO); Liang, 2020), whereas the median age for SARS patients in China was 33 years old (Cao et al., 2011; Feng et al., 2009). It is tempting to speculate that the opposite susceptibility of old and young people to SARS-CoV and SARS-CoV-2 may relate to the differential usage of ANPEP and ACE2 receptors by these two closely-related coronaviruses. However, our analysis is limited by a small sample size and many possible confounders such as gender, smoking status, and other genetic and environmental factors, which could not be controlled for.

### Digestive system: infection hotspot?

Our interrogation of SCARF expression in the major organs of the digestive system (GI tract, liver, pancreas, gallbladder) provides confirmation of previous findings and yields novel observations. Consistent with several studies (Hikmet et al., 2020; Lee et al., 2020; Sungnak et al., 2020; Ziegler et al., 2020), we found that the small intestine is one of the ‘hottest’ tissues for co-expression of *TMPRSS2* with *ACE2*, but also for *DPP4* and *ANPEP* as reported previously in two other studies (Liang et al., 2020; Venkatakrishnan et al., 2020). Within the small intestine, we found that the jejunum is where highest expression of these factors is achieved. This is in slight deviation from Ziegler et al. who suggested that the ileum had the maximum expression of *ACE2* (Ziegler et al., 2020). Regardless of which section of the small intestine, both studies converge on the finding that expression of these factors is largely driven by enterocytes and their progenitors, which line the inner surface of the intestine and are therefore directly exposed to food and pathogens.

Goblet cells represent another cell type commonly found in the digestive system that we also predict to be permissive for SARS-CoV-2 entry. These are epithelial cells found in the airway, intestine, and colon that specialize in mucosal secretion. We found that goblet cells have some of the highest level of co-expression of *TMPRSS2* with one or several receptors including *ACE2, ANPEP, DPP4* and *CD147*/BSG. The same cell type also displayed high levels of Cathepsin B and L, which may also facilitate SARS-CoV-2 infection of the digestive system. Goblet cells within the nasal epithelium have also been identified as potentially vulnerable to SARS-CoV-2 (Sungnak et al., 2020). Overall, our analysis of the digestive system is concordant with several other studies pointing at the lining of the GI tract as common site of SARS-CoV-2 infection (Lamers et al., 2020). This could explain the digestive symptoms (e.g. diarrhea) presented in COVID-19 cases (Wang et al., 2020a) as well as the detection of viral shedding in feces (Xu et al., 2020a). If so, fecal-oral transmission of SARS-CoV-2 may be plausible, but it remains to be rigorously investigated.

### Concluding remarks

Overall, this study provides a valuable resource for future studies of the basic biology of SARS-CoV-2 and other coronaviruses as well as clinical investigations of the pathology and treatment of COVID-19. We also established an open-access, web-based interface, dubbed SCARFace, allowing any user to explore the expression of any SCARF (and any other human RefSeq gene) at single-cell resolution within any of the scRNA-seq dataset analyzed here. Our finding that SCARF expression is generally well conserved across primate species, along with high level of sequence conservation of the ACE2 interface with the Spike protein (Damas et al., 2020; Melin et al., 2020), suggests that non-human primates are adequate models for the study of SARS-CoV-2 and the development of therapeutic interventions, including vaccines. It would be desirable to obtain single-cell resolution of SCARF expression in a broader range of tissues and species, including non-primate models such as hamsters and ferrets. While our survey of SCARF expression across human embryonic and adult tissues is the broadest of its kind, it remains limited by the constraints and shortcomings of scRNA-seq. These include the lack or under-representation of certain cell types that are rare or undetected due to low sequencing depth, isolation biases, or statistical cutoffs. Likewise, the expression level of any given gene may be underestimated due to dropout effects. Hence, we strongly recommend interpreting our results with caution, especially negative results. Furthermore, RNA expression levels are imprecisely reflective of protein abundance and our observations need to be corroborated by approaches quantifying protein expression *in situ*. Lastly, and perhaps most critically, SCARF expression within and between individuals is bound to be heavily modulated by genetic and environmental factors, including infection by SARS-CoV-2 and other pathogens. Such variables may drastically shift the expression patterns we observe in healthy tissues from a limited number of donors. In fact, several SCARFs surveyed here such as IFITM and LY6E restriction factors (Jia et al., 2012; Mar et al., 2018), and apparently ACE2 itself (Ziegler et al., 2020), are known to be modulated by infection and the innate immune response. Our data still provide a valuable baseline to evaluate how SCARF expression may be altered during the course of SARS-CoV-2 infection. Because we survey host factors associated with a range of zoonotic coronaviruses, this study may also prove a useful resource in the context of other eventual outbreaks.

## ACKNOWLEDGEMENTS

We would like to thank members of the Feschotte Lab and of the Cornell Virology community, as well as Ankit Arora for helpful advice and discussions. M.S. is supported by a Presidential Postdoctoral Fellowship from Cornell University. V.B. is supported by a Career Development Fellowship at DZNE Tuebingen. Work on host-virus interactions in the Feschotte lab is funded by R35 GM122550 from the National Institutes of Health. Figure 1 was created with BioRender.com.

## AUTHOR CONTRIBUTIONS

C.F., M.S. and V.B. conceived the study. C.F. supervised the project and wrote the manuscript. M.S. and V.B. assisted in writing, carried out all data analysis, and constructed the web interface ‘*SCARFace’*.

## SUPPLEMENTAL FIGURE TITLES AND LEGENDS

**Figure S1:**
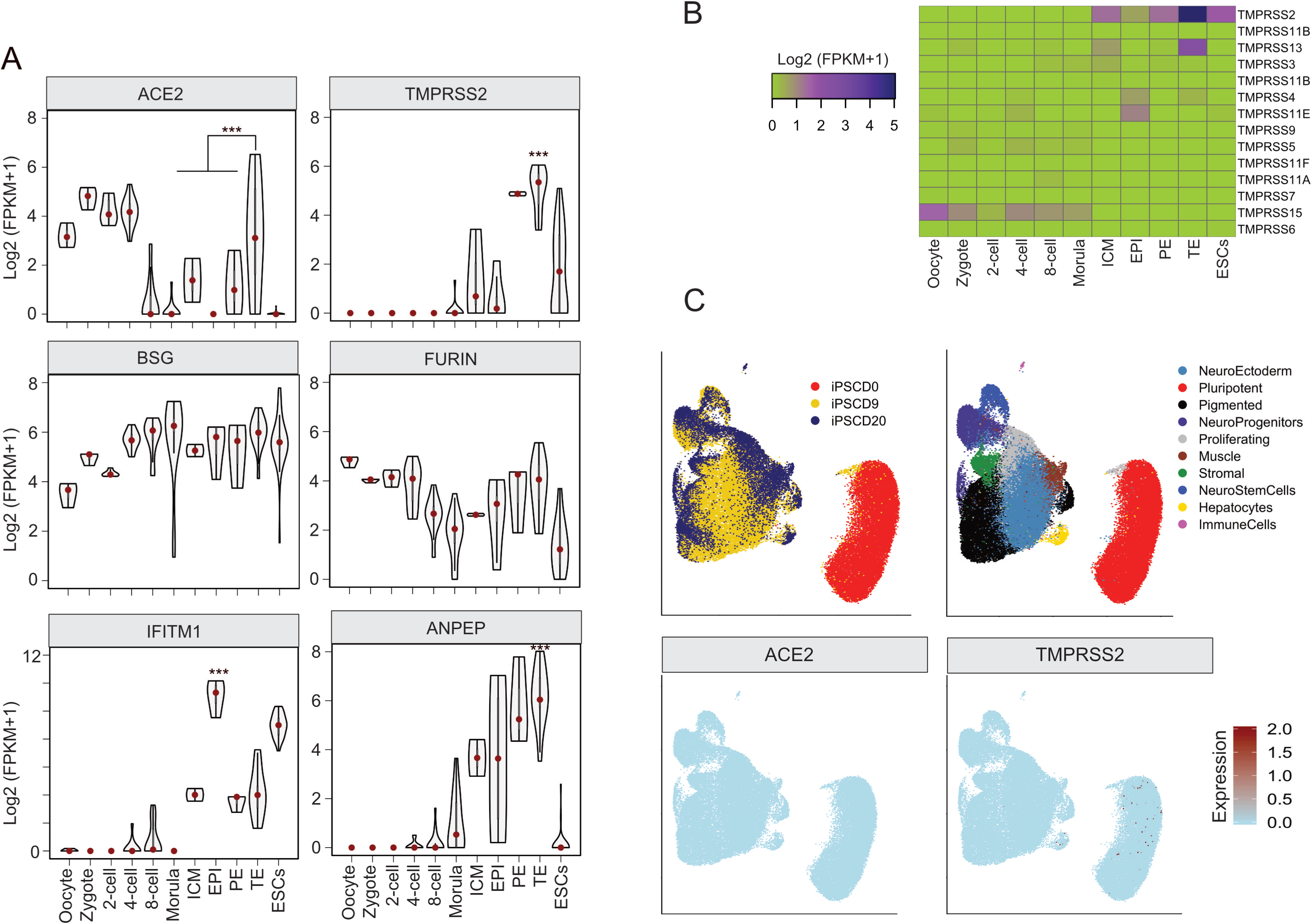
Expression of SCARFs during the differentiation of Pluripotent cells. A. Violin plots show the Log2 normalized expression of ACE2, TMPRSS2, ANPEP, BSG, FURIN, and IFITM1 in the preimplantation embryos of humans from the analysis of single-cell scRNA-seq datasets. B. Heatmap visualization of transcript expression [log2 RPKM] values of TMPRSS genes for each stage of human preimplantation embryos. The Colour scheme: low expressed (green), to highly expressed (purple). C. Upper panel: UMAP bi-plot shows the clustering of ∼ 60,000 scRNA-seq datasets from human induced pluripotent cell differentiation till day 9 and day 20. Lower panel: Two feature plots based on the UMAP plot show the expression of CoV receptor (ACE2) and protease (TMPRSS2).

**Figure S2:**
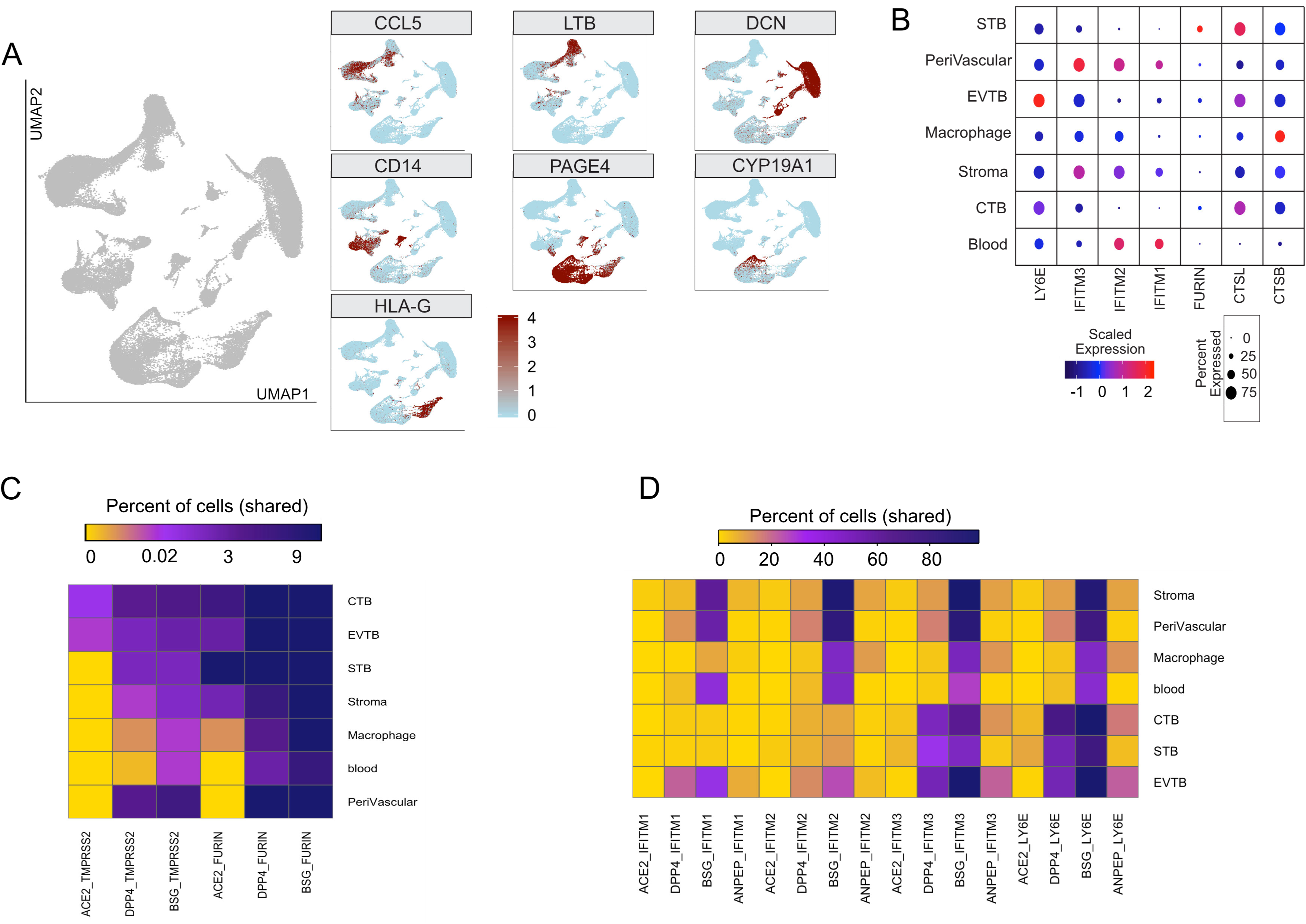
Placental cells could be vulnerable to SARS-CoV-2 infection. A. Multiple UMAP plots show the clusters (Figure 2B) can be distinguished by the strong expression of known markers CTB (PAGE4), STB (CYP19A1), EVTB (HLA-G), Decidual cells (DCN), Macrophages (CD14), and Immune cells (CCL5 and LTB). B. Dot plot illustrates the intensity and density of SARS-CoV proteases, and restriction factor gene expression between the cell-types shown in the previous figure. Size and color of dots are as in Figure 2C, 3A, C and 4B. C. Heatmap showing the fractions of receptors and proteases double-positive cells that are enriched in any placental lineages. The Colour scheme is based on distribution from a low (< 0.02%, gold), medium (0.02-3% purple), and a high fraction (3-8, and above %, blue) of double-positive cells enriched in a given cell type. D. Heatmap showing the fractions of receptors and restriction factors double-positive cells. Colour scheme is based on distribution from low fraction (< 20 %, gold), medium fraction (20-60% purple), high fraction (> 60 %, blue) of double-positive cells enriched in a particular cell type.

**Figure S3:**
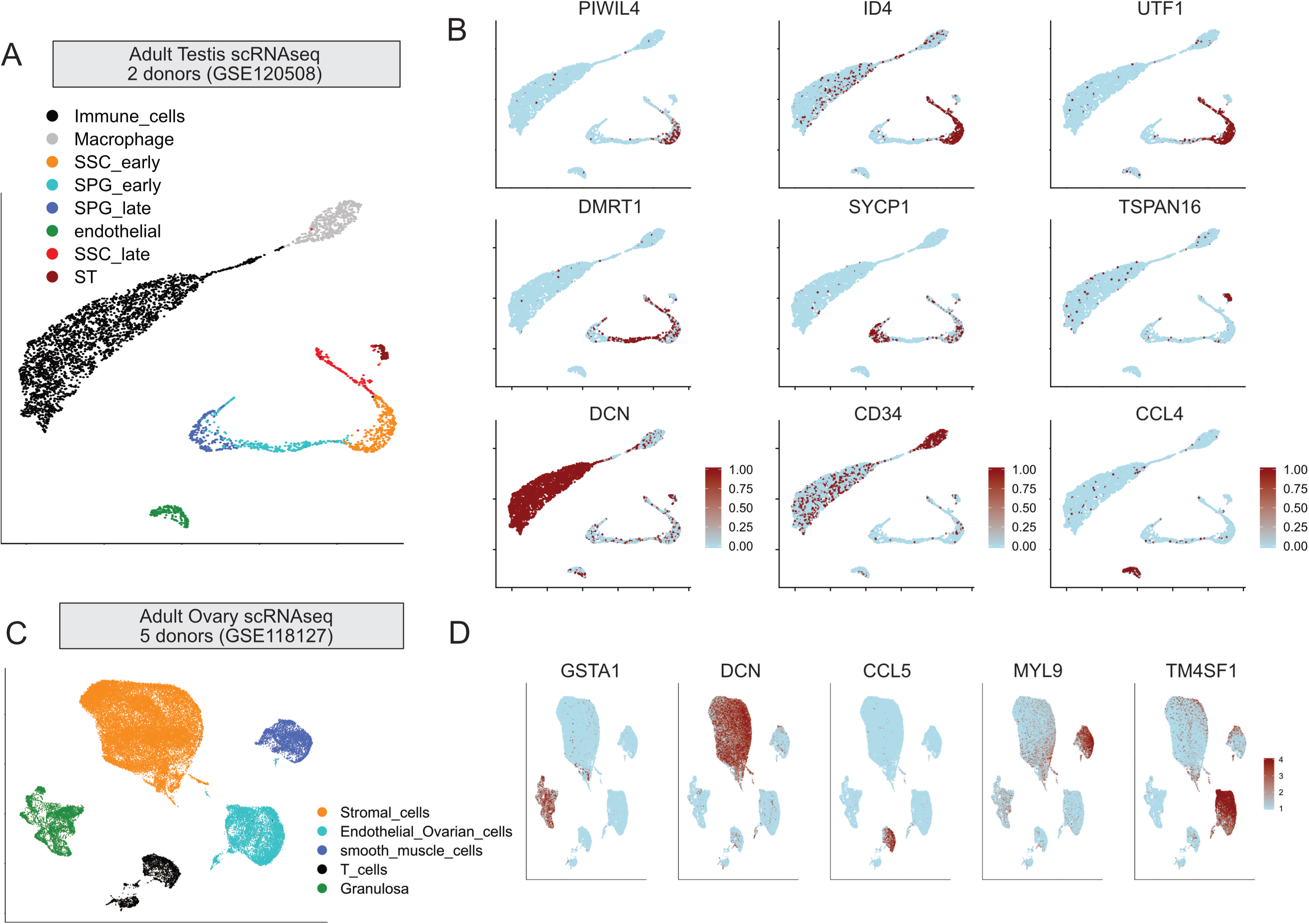
Single cellular clustering of the adult testis and ovary from 2 and 5 individuals respectively. A. Left panel: UMAP biplot of the adult human testis (N=2). The cells in each cluster are spermatogonia stem cell (SSC), spermatogonia progenitors (SPG), spermatid (ST), endothelial cells, and macrophages. Right panel: The feature plots show the expression of markers, taken from the original article in individual cells. B. Left panel: UMAP biplot of the adult ovary (N=5). The cells in each cluster are Ovarian granulosa, stromal cells, endothelial cells, smooth muscle cells, and T cells. Right panel: The feature plots show the expression of markers (taken from the original article in individual cells.

**Figure S4:**
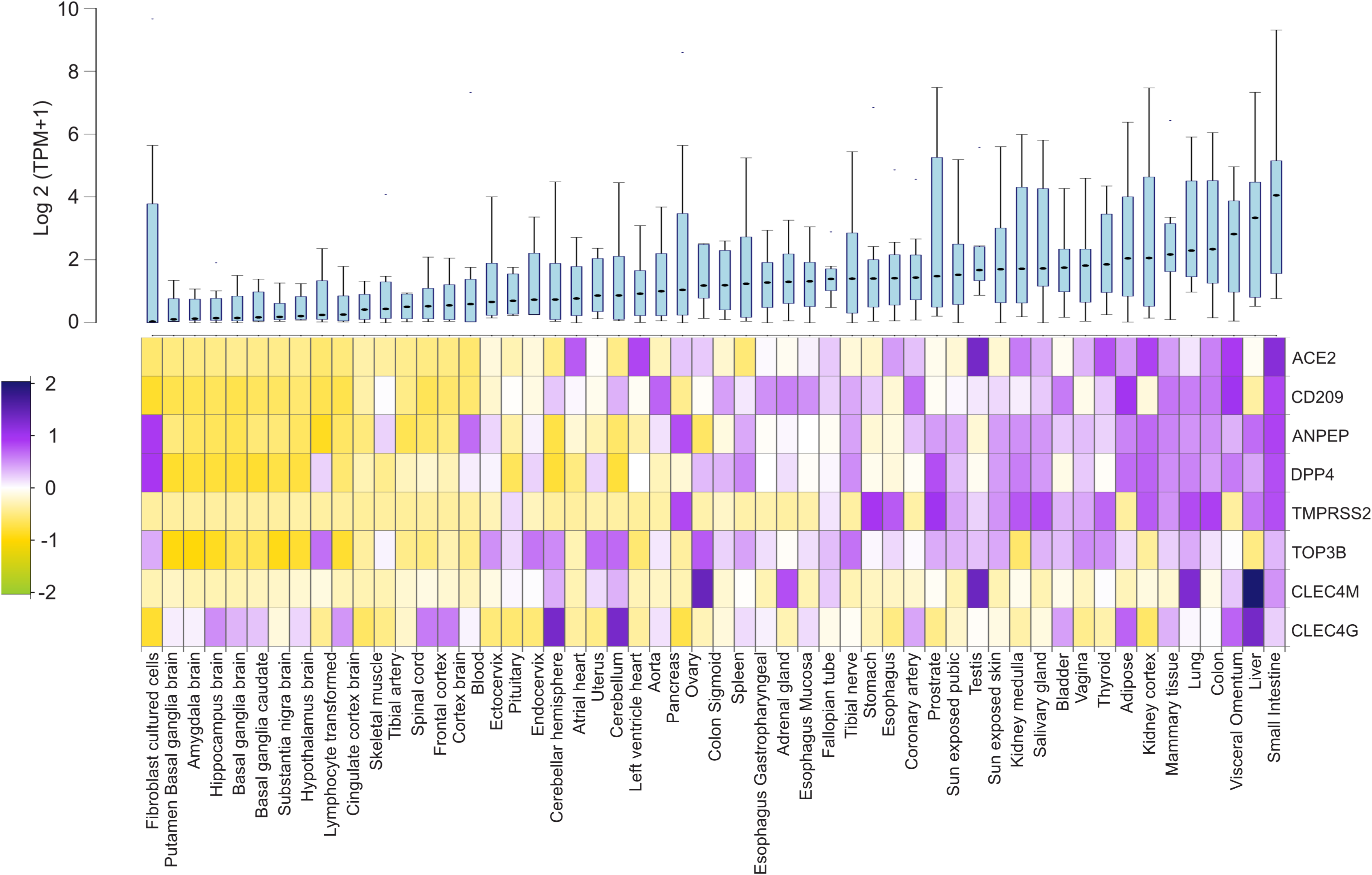
Expression of Receptors from the bulk RNA-seq (GTEx) The boxplot shows the distribution of overall expression of the shown CoV entry factors in the distinct post-mortem tissues from GTEx datasets. The heatmap beneath the boxplot displays scaled expression of individual genes, the upregulation of shown factors in digestive, reproductive, endocrine, and respiratory organ systems.

**Figure S5:**
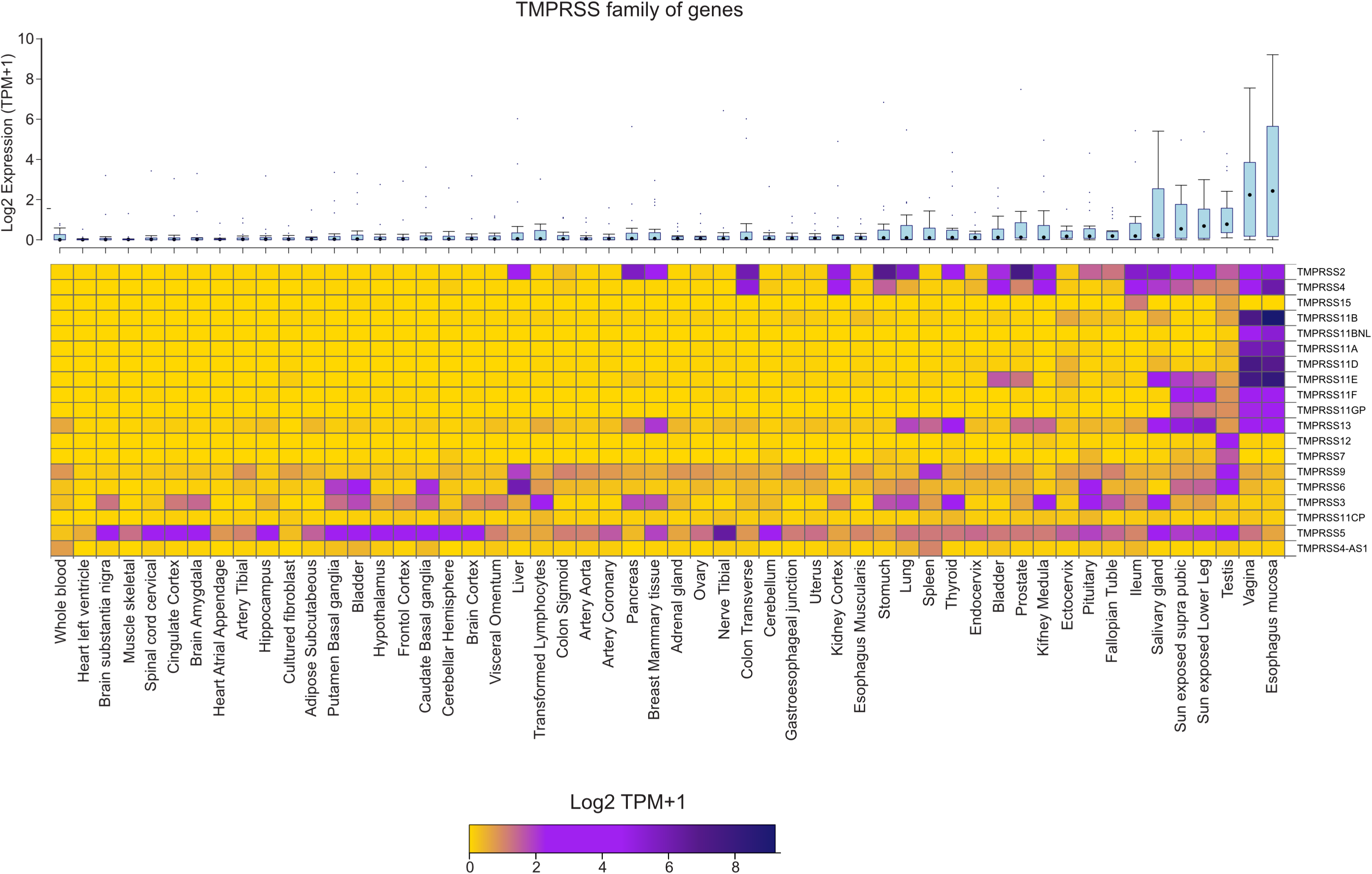
Expression of TMPRSS family of Proteases from the bulk RNA-seq (GTEx) The boxplot shows the distribution of the overall expression of TMPRSS family genes in post-mortem tissues from GTEx datasets. The heatmap beneath the boxplot displays the expression (log2 (TPM+1)) of individual genes.

**Figure S6:**
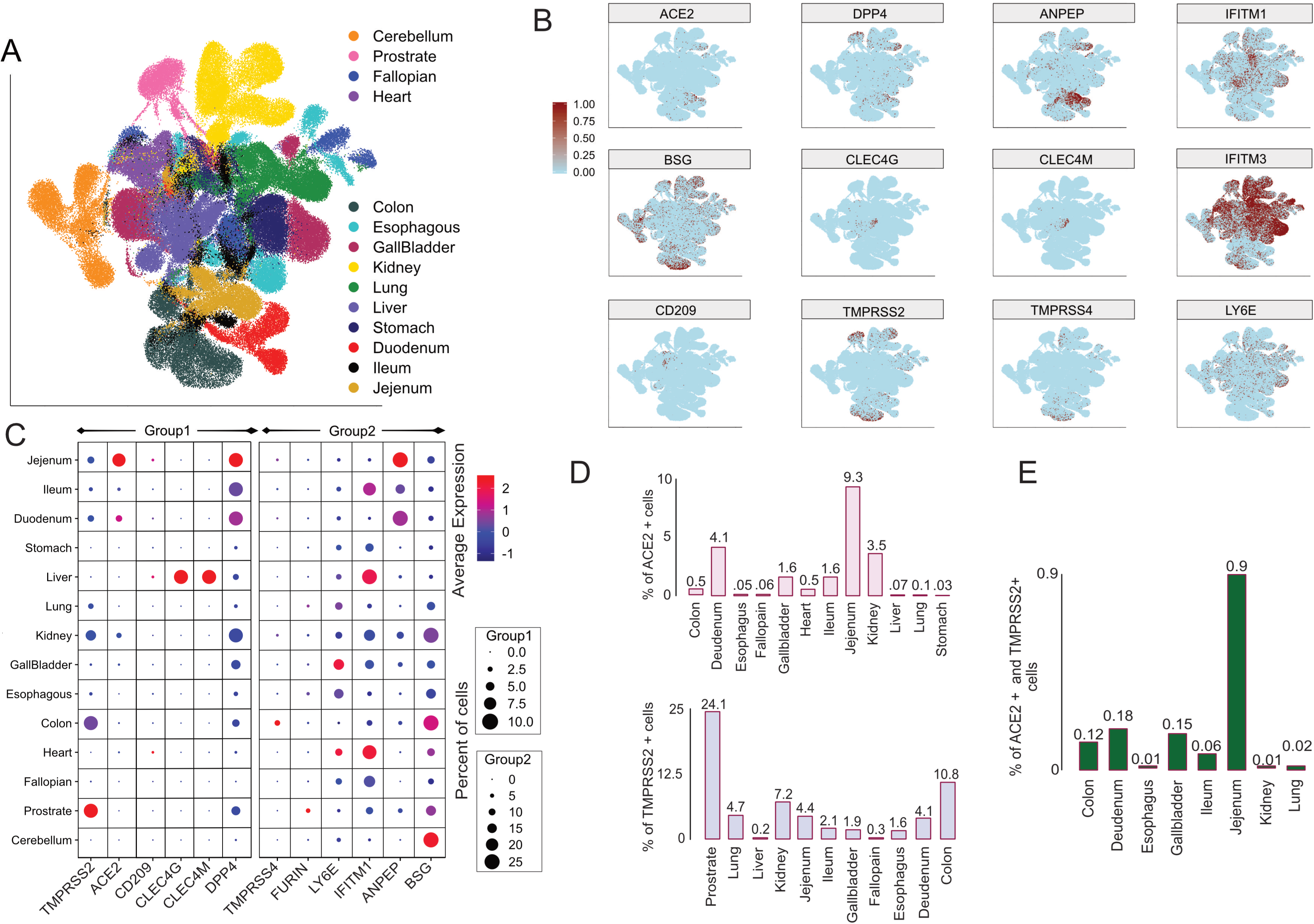
Expression of SCARFs from the HCL. A. UMAP biplot of ∼ 200,000 cells from 14 distinct adult tissues visualized through unsupervised clustering. Every dot is a single cell corresponding to a labeled tissue coded by different colors (See also Figure 4). B. Multiple feature plots demonstrating the single cellular expression of selected SCARFs (all seven entry receptors, proteases i.e., TMPRSS2/4, and restriction factors i.e., *IFITM1/2/3 and LY6E*) across the tissues shown in the previous UMAP plot. C. Dot plot showing the abundance and intensity of transcript expression of selected SCARFs (all seven entry receptors, proteases i.e., *TMPRSS2/4 and FURIN,* and restriction factors i.e., *IFITM1 and LY6E*) across the analyzed 14 tissues. See legend of Figure 2C for a description of dot plot display. D. Bar plots are showing the percent of cells expressing ACE2 (upper panel) and TMPRSS2 (lower panel) in each tissue. Only those tissues are shown that have any cell expressing the respective genes. E. The bar plot represents the percent of *ACE+ TMPRSS2+* cells in each tissue. Only those tissues are shown that have any cell co-expressing the *ACE2* and *TMPRSS2*.

**Figure S7:**
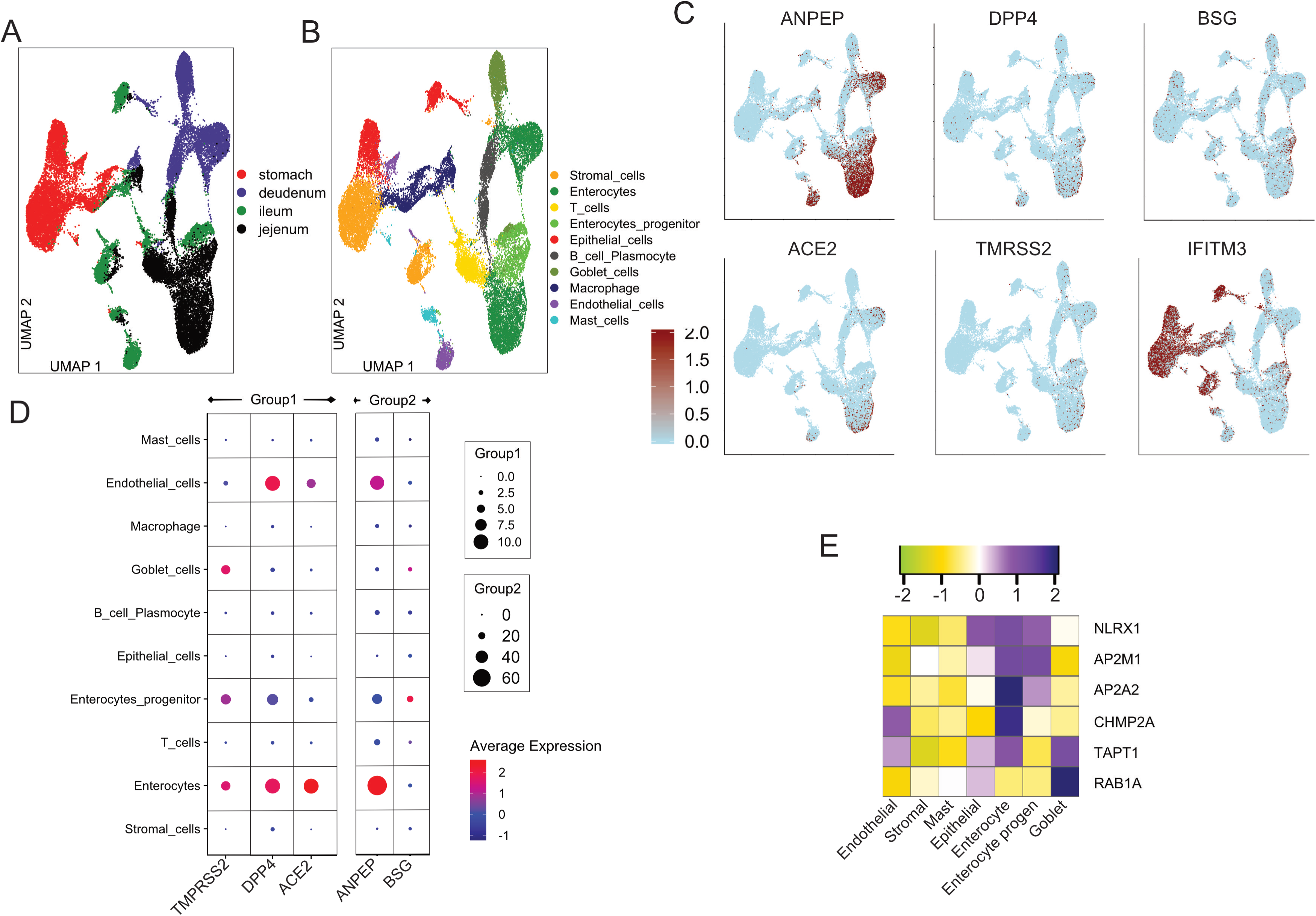
Expression of SCARFs from the digestive tract. A. UMAP biplot of cells from digestive tract tissues (stomach, ileum, duodenum, and jejunum) visualized through unsupervised clustering. Every dot is a single cell corresponding to a labeled tissue coded by different colors. B. Visualization of 10 cell types in the digestive tract tissues (See Figure S7A) obtained using unsupervised clustering. Clusters were annotated using the known markers from the original article. C. Multiple feature plots demonstrating the single cellular expression of selected SCARFs (4 CoV receptors that are broadly expressed in digestive tract i.e. *ACE2, ANPEP, DPP4, and BSG*, a protease i.e., *TMPRSS2*, and a restriction factors i.e. *IFITM1*) across the dataset visualized in the previous UMAP plot. D. Dot plot showing the transcript expression of CoV receptors demonstrated in the previous figure, and *TMPRSS2* across the analyzed ten cell types. See legend of Figure 2C for a description of dot plot display. E. Heatmap is displaying the scaled normalized expression (averaged for each type) of the intracellular facilitating factors enlisted in SCARFs (Table S2). Here, we only show those factors that are among the most variable genes across the dataset.

**Figure S8:**
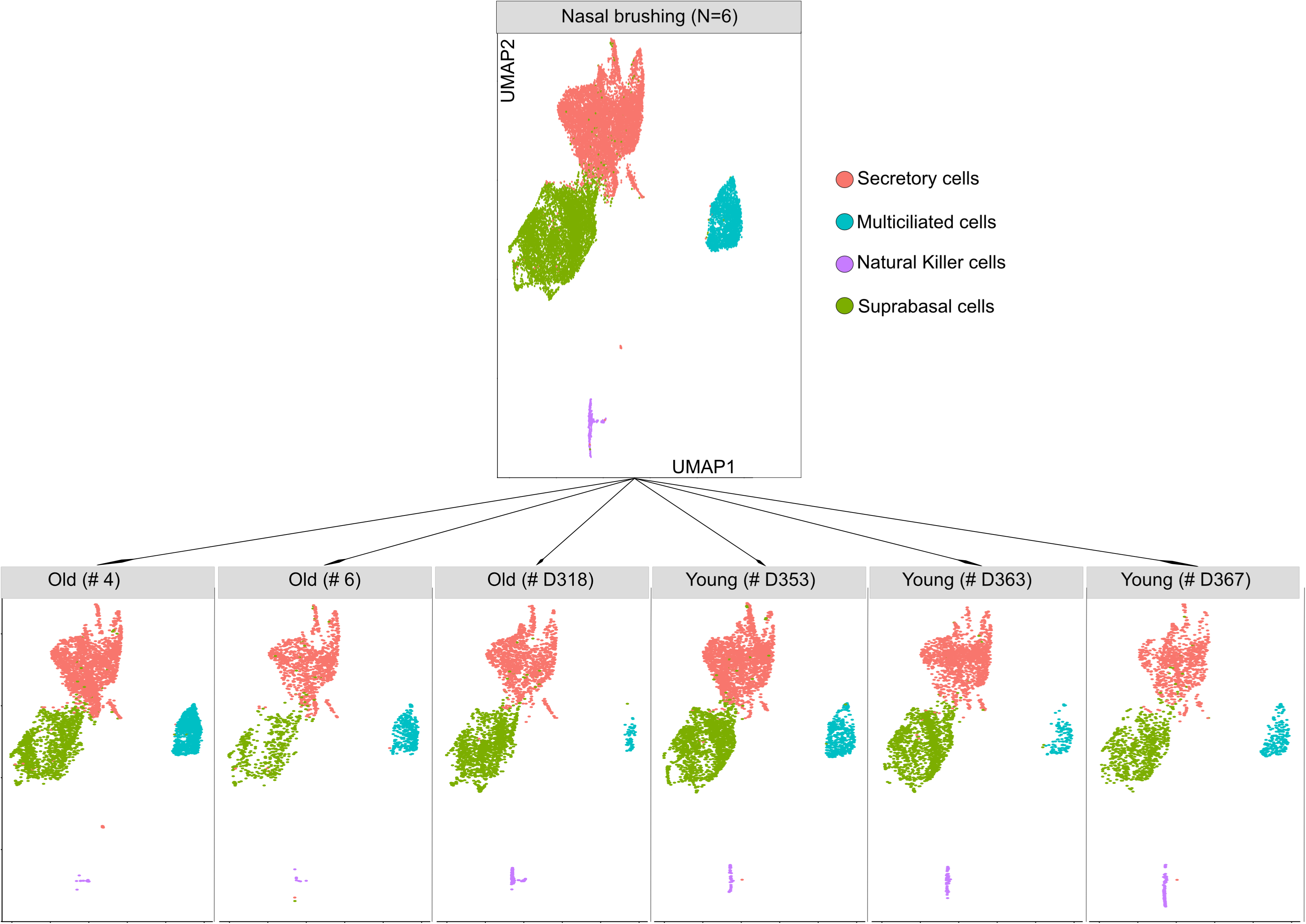
Integration of Nasal epithelial scRNA-seq datasets from 3 studies. Upper panel: Integrative analysis of 6 scRNA-seq datasets obtained from 3 studies reveals four specific clusters, representing the major nasal epithelial cell types i.e., ciliated, secretory, suprabasal, and natural killer (NK) cells. Lower Panel: UMAP plots of all six samples separately showing the contribution of each sample to the clusters above. Note: 3 samples were obtained from older (age, 50-59), and the rest 3 were from younger individuals (age, 24-30).

**Figure S9:**
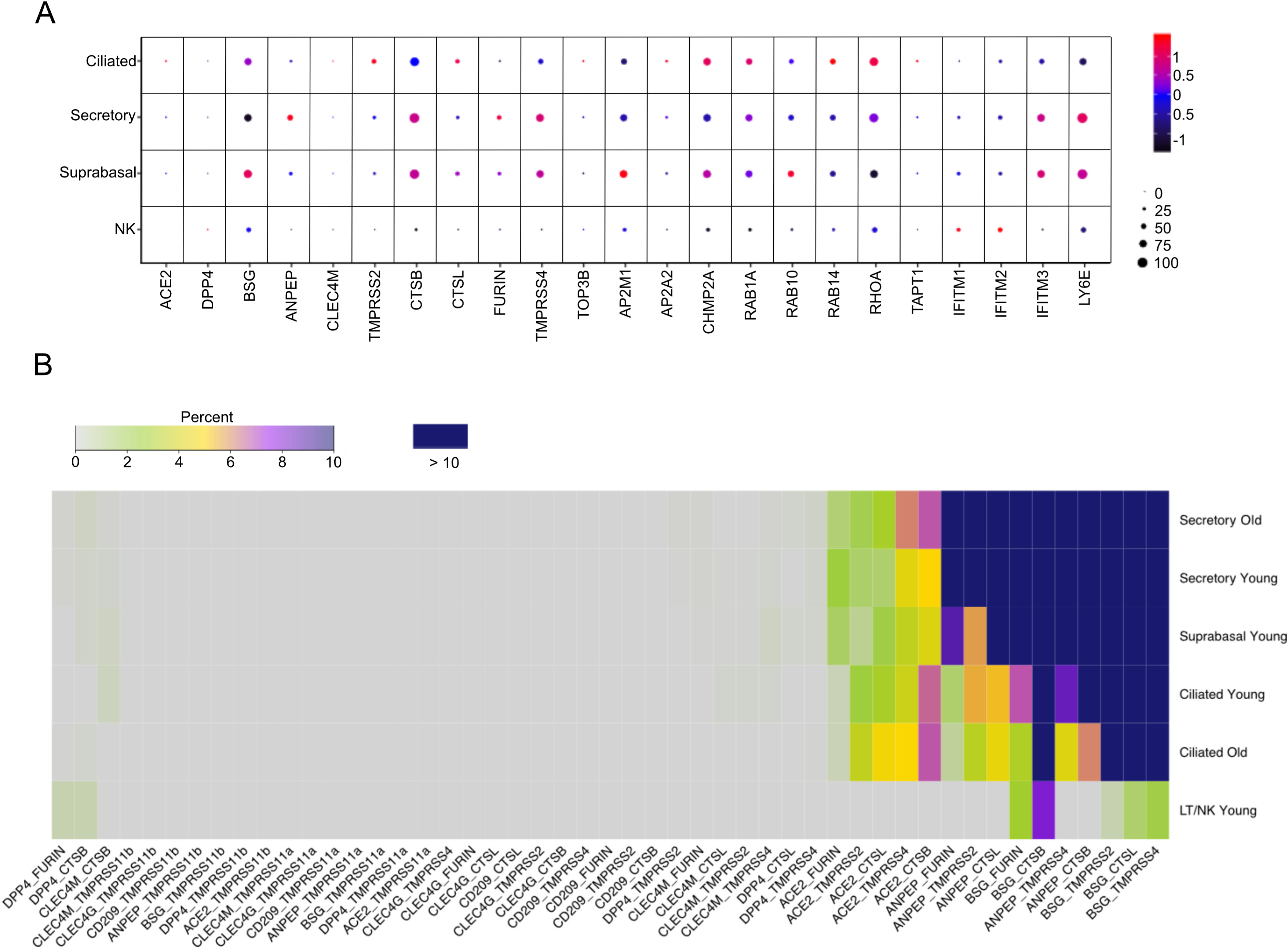
Expression of SCARFs in nasal epithelial cells. A. Dot plot showing the transcript expression of SCARFs across the 4 major cell types in nasal epithelial cells. B. Heatmap is showing the double-positive cells for CoV receptors and proteases in the mentioned cell types from older and younger individuals.

## SUPPLEMENTAL TABLES

**Table S1:** List of studies profiling the expression of SARS-CoV-2 entry factors in human tissues

**Table S2:** List of SARS-CoV-2 and coronavirus-associated receptors and factors (SCARFs)

**Table S3:** Overview of the datasets used in the study

**Table S4:** Percentage of positive, double-positive cells in maternal fetal interface, testis and somatic tissues

**Table S5:** Markers identified from scRNA-seq data

**Table S6:** Percentage of positive/double-positive cells in nasal epithelial cells

**Table S7:** Differentially expression genes between ciliated and secretory nasal cells

## METHODS

### Pre-implantation embryos

Single-cell (sc) RNA-seq datasets from the pre-implantation stages of development were downloaded in a raw format from ((Yan et al., 2013), GSE36552). RNAseq reads with MAP quality score < 30 were removed. Resulting reads were mapped to the human genome (hg19) using STAR (https://github.com/alexdobin/STAR) with defined settings, i.e. --*alignIntronMin 20 --alignIntronMax 1000000 --chimSegmentMin 15 --chimJunctionOverhangMin 15 --outFilterMultimapNmax 20*, and only uniquely mapped reads were considered for the calculation of expression. RPKM was calculated using *bamutils* (http://ngsutils.org/modules/bamutils/count/) for individual genes annotated in the human RefSeq database. Note: Here, we used the expression matrix generated for a previously published work (Izsvák et al., 2016), using the above mentioned pipeline.

### Maternal-fetal interface

We obtained the processed expression matrix (counts) from ((Vento-Tormo et al., 2018), E-MTAB-6701) for ∼ 70,000 single cells representing the maternal-fetal interface. We then used Seurat (v3.1.1) (https://github.com/satija.lab/seurat) within the R environment (v3.6.0) for the processing the dataset. We kept the cells with minimum and maximum of 1,000 and 5,000 genes expressed (≥1 count), respectively. Moreover, cells with more than 5% of counts on mitochondrial genes were filtered out. After filtering, there were 64,782 cells. The data normalization was achieved by scaling it with the factor 10,000 followed by natural-log transformation. Clustering was performed using the “*FindClusters*” function with default parameters, except the resolution was set to 0.1. We used the first 20 Principle Component (PC) dimensions in the construction of the shared-nearest neighbor (SNN) graph to generate 2-dimensional embeddings for data visualization using UMAP. Cell type assignment was performed based on the annotations provided by the original publication, albeit we grouped the clusters into broader lineages. T-cell, B-cell, Dendritic, NK-cells, and Monocytes were categorized into “blood,” all decidual cells, except perivascular cells, were annotated as “stroma.” Fetal lineages were grouped into the known groups as ExtravillousTrophoblast (“EVTB”), CytoTrophoblast (“CTB”), and Syncytiotrophoblast (“STB”) cells. All the given annotations were further confirmed by their respective markers (Figure S2).

### Adult tissues

We mined the scRNA-seq of 2 tissue samples from adult Testis ((Sohni et al., 2019), GSE124263), 31 tissue samples from 5 ovaries ((Wagner et al., 2020), GSE118127), 29 samples of 14 adult tissues from Human Cell Landscape (HCL) ((Han et al., 2020), GSE134355) in the form of raw counts. To avoid the cross-platform batch biasedness, we independently processed the samples taken from different studies. Datasets corresponding to the same tissue were merged into one before the downstream processing. We scaled and normalized the datasets Seurat (v3.1.1) (https://github.com/satija.lab/seurat) within the R environment (v3.6.0) as described in the previous section. We used a similar pipeline for the downstream analysis, too, except that for merged HCL samples. We fed the first 50 PCs as input to cluster and visualize the single cells using SNN graphs and UMAP methods. The top marker genes distinguishing the cell types were calculated using *the “FindAllMarkers”* function implemented in Seurat, (adjusted *p-value* < 0.01 and log(fold-change) > 0.25) using the Wilcoxon Rank Sum test. We annotated the cell types using the markers obtained in this study and cross-referenced with the original article.

### Nasal epithelium

In order to compare young and old nasal tissue cells, we defined samples with age ≤ 30 as young and with age ≥ 50 as old. We found no study that has healthy old and young nasal samples within the same experiment. Therefore, six samples have been taken from three studies ((Deprez et al., 2019; Garcıá et al., 2019; Vieira Braga et al., 2019), Table S3). Note that cell annotations were used as provided by the original publications except for the sample D318 (raw count matrix was available). Unless mentioned specifically, broad annotation terms as ciliated and secretory has been used for the sake of consistency.

#### Processing of samples “4”, “6”, “D353”, “D363” and “D367”

Normalized counts and cell-type annotations provided by the original publications were used. “CellType” annotations with ≥ 100 cells were considered, giving “Ciliated_2” (n=1,513), “Goblet_2” (n=1,463) and “Goblet_1” (n=4,017) for old samples “4” and “6”, and “LT/NK” (n=185), “Multiciliated_N” (n=855), “Secretory_N” (n=7,138) and “Suprabasal_N” (n=1,640) for young samples “D353”, “D363” and “D367”.

#### Processing of sample D318 raw matrix

Firstly, cellranger (v3.1.0) *reanalyze* was used to generate filtered matrix of top 5,000 cells (https://github.com/10XGenomics/cellranger). Next, we used Scrublet (v0.2.1) (Wolock et al., 2019) for identifying doublets with expected doublet rate of 0.03. 58 cells were discarded with scores higher than 0.2. SoupX (v1.2.2) (Young and Behjati, 2018) was used to subtract the ambient RNA profiles from the real expression values. Finally, we used Seurat (v3.1.1) within the R environment (v3.6.0) for filtering, normalization and cell-type identification for sample D318. The following data processing was done: (1) Filtering. We kept the cells with minimum and maximum of 1,000 and 5,000 genes expressed (≥1 count), respectively. Moreover, cells with more than 10% of counts on mitochondrial genes were filtered out. After filtering, there were 2,987 cells. (2) Data normalization. Gene UMI counts for each cell were divided by the total number of counts in that cell and multiplied by 10,000. These values were then natural-log transformed. (3) Cell-type identification. Integration of sample D318 scRNA-seq data with remaining samples was performed using top 2000 variable features. Clustering was performed using “*FindClusters*” function with default parameters except resolution was set to 0.1 and first 30 PCA dimensions were used in the construction of the shared-nearest neighbor (SNN) graph and to generate 2-dimensional embeddings for data visualization using UMAP. Cell types were assigned based on the annotations provided by the original publication of samples “D353”, “D363” and “D367”, giving “LT/NK” (n=110), “Multiciliated_N” (n=62), “Secretory_N” (n=1,354) and “Suprabasal_N” (n=1,461).

#### Percentage analysis

For each sample, the number of positive cells for a gene was calculated when they had a count higher than 0. The numbers were added separately for young and old samples. Percentage was calculated for each cell type. P-value was calculated between the percentage of positive cells in young samples and old samples using one-sided fisher’s exact test.

#### Differential expression analysis

We used the “*FindAllMarkers*” function with default parameters except for a minimum percentage set to 5%. Default cutoffs were used to identify significant DE genes with log FC of |0.25| and adjusted p-value of less than 0.01. Genes below these cutoffs are shown in volcano plots for visualization purpose only.

### Cross-species analysis

Trimmed mean of M values (TMM) normalized cross-species Counts per million (CPM) values were imported in R and variable features were identified using “*FindVariableFeatures*” function implemented in the Seurat package using *mean.var.plot (mvp)* as a selection method. Clustering was performed using “*FindClusters*” function with default parameters except resolution was set to 1 and first 10 PCA dimensions were used in the construction of the shared-nearest neighbor (SNN) graph and to generate 2-dimensional embeddings for data visualization using UMAP.

## REFERENCES

Aguet, F., Brown, A.A., Castel, S.E., Davis, J.R., He, Y., Jo, B., Mohammadi, P., Park, Y.S., Parsana, P., Segrè, A. V., et al. (2017). Genetic effects on gene expression across human tissues. Nature 550, 204–213.

Aguiar, J.A., Tremblay, B.J.-M., Mansfield, M.J., Woody, O., Lobb, B., Banerjee, A., Chandiramohan, A., Tiessen, N., Dvorkin-Gheva, A., Revill, S., et al. (2020). Gene expression and in situ protein profiling of candidate SARS-CoV-2 receptors in human airway epithelial cells and lung tissue. BioRxiv.

Aylward, Bruce (WHO); Liang, W. (PRC) (2020). Report of the WHO-China Joint Mission on Coronavirus Disease 2019 (COVID-19).

Baud, D., Greub, G., Frave, G., Gengler, C., Jaton, K., Dubruc, E., and Pomar, L. (2020). Second-Trimester Miscarriage in a Pregnant Woman With SARS-CoV-2 Infection. JAMA - J. Am. Med. Assoc.

Bertram, S., Glowacka, I., Müller, M.A., Lavender, H., Gnirss, K., Nehlmeier, I., Niemeyer, D., He, Y., Simmons, G., Drosten, C., et al. (2011). Cleavage and activation of the severe acute respiratory syndrome coronavirus spike protein by human airway trypsin-like protease. J. Virol. 85, 13363–13372.

Blake, L.E., Roux, J., Hernando-Herraez, I., Banovich, N.E., Perez, R.G., Hsiao, C.J., Eres, I., Cuevas, C., Marques-Bonet, T., and Gilad, Y. (2020). A comparison of gene expression and DNA methylation patterns across tissues and species. Genome Res. 30, 250–262.

Blanco-Melo, D., Nilsson-Payant, B.E., Liu, W.-C., Uhl, S., Hoagland, D., Møller, R., Jordan, T.X., Oishi, K., Panis, M., Sachs, D., et al. (2020). Imbalanced host response to SARS-CoV-2 drives development of COVID-19. Cell.

Cao, W.C., de Vlas, S.J., and Richardus, J.H. (2011). The severe acute respiratory syndrome epidemic in mainland China dissected. Infect. Dis. Rep. 3, 3–6.

Chen, H., Guo, J., Wang, C., Luo, F., Yu, X., Zhang, W., Li, J., Zhao, D., Xu, D., Gong, Q., et al. (2020a). Clinical characteristics and intrauterine vertical transmission potential of COVID-19 infection in nine pregnant women: a retrospective review of medical records. Lancet 395, 809–815.

Chen, R., Wang, K., Yu, J., Chen, Z., Wen, C., and Xu, Z. (2020b). The spatial and cell-type distribution of SARS-CoV-2 receptor ACE2 in human and mouse brain. BioRxiv 2020.04.07.030650.

Chen, Z., Mi, L., Xu, J., Yu, J., Wang, X., Jiang, J., Xing, J., Shang, P., Qian, A., Li, Y., et al. (2005). Function of HAb18G/CD147 in Invasion of Host Cells by Severe Acute Respiratory Syndrome Coronavirus. J. Infect. Dis. 191, 755–760.

Cheng, Y., Luo, R., Wang, K., Zhang, M., Wang, Z., Dong, L., Li, J., Yao, Y., Ge, S., and Xu, G. (2020). Kidney impairment is associated with in-hospital death of COVID-19 patients. MedRxiv 2020.02.18.20023242.

Chow, R.D., and Chen, S. (2020). The aging transcriptome and cellular landscape of the human lung in relation to SARS-CoV-2. BioRxiv 2020.04.07.030684.

Corman, V.M., Muth, D., Niemeyer, D., and Drosten, C. (2018). Hosts and Sources of Endemic Human Coronaviruses. In Advances in Virus Research, p.

Cui, P., Chen, Z., Wang, T., Dai, J., Zhang, J., Ding, T., Jiang, J., Liu, J., Zhang, C., Shan, W., et al. (2020). Clinical features and sexual transmission potential of SARS-CoV-2 infected female patients: a descriptive study in Wuhan, China. MedRxiv 2020.02.26.20028225.

Damas, J., Hughes, G.M., Keough, K.C., Painter, C.A., Persky, N.S., Corbo, M., Hiller, M., Koepfli, K.-P., Pfenning, A.R., Zhao, H., et al. (2020). Broad Host Range of SARS-CoV-2 Predicted by Comparative and Structural Analysis of ACE2 in Vertebrates. BioRxiv 2020.04.16.045302.

Deprez, M., Zaragosi, L.-E., Truchi, M., Garcia, S.R., Arguel, M.-J., Lebrigand, K., Paquet, A., Pee’r, D., Marquette, C.-H., Leroy, S., et al. (2019). A single-cell atlas of the human healthy airways. BioRxiv.

Diao, B., Feng, Z., Wang, C., Wang, H., Liu, L., Wang, C., Wang, R., Liu, Y., Liu, Y., Wang, G., et al. (2020). Human Kidney is a Target for Novel Severe Acute Respiratory Syndrome Coronavirus 2 (SARS-CoV-2) Infection. MedRxiv 2, 2020.03.04.20031120.

Ding, Y., Wang, H., Shen, H., Li, Z., Geng, J., Han, H., Cai, J., Li, X., Kang, W., Weng, D., et al. (2003). The clinical pathology of severe acute respiratory syndrome (SARS): A report from China. J. Pathol.

Domínguez-Soto, A., Aragoneses-Fenoll, L., Gómez-Aguado, F., Corcuera, M.T., Clária, J., García-Monzón, C., Bustos, M., and Corbí, A.L. (2009). The pathogen receptor liver and lymph node sinusoidal endotelial cell C-type lectin is expressed in human Kupffer cells and regulated by PU.1. Hepatology 49, 287–296.

Fanelli, V., Fiorentino, M., Cantaluppi, V., Gesualdo, L., Stallone, G., Ronco, C., and Castellano, G. (2020). Acute kidney injury in SARS-CoV-2 infected patients. Crit. Care 24.

De Felice, F.G., Tovar-Moll, F., Moll, J., Munoz, D.P., and Ferreira, S.T. (2020). Severe Acute Respiratory Syndrome Coronavirus 2 (SARS-CoV-2) and the Central Nervous System. Trends Neurosci. 0.

Feng, D., De Vlas, S.J., Fang, L.Q., Han, X.N., Zhao, W.J., Sheng, S., Yang, H., Jia, Z.W., Richardus, J.H., and Cao, W.C. (2009). The SARS epidemic in mainland China: Bringing together all epidemiological data. Trop. Med. Int. Heal. 14, 4–13.

Frieman, M., and Baric, R. (2008). Mechanisms of Severe Acute Respiratory Syndrome Pathogenesis and Innate Immunomodulation. Microbiol. Mol. Biol. Rev. 72, 672–685.

Gao, Q.Y., Chen, Y.X., and Fang, J.Y. (2020). 2019 Novel coronavirus infection and gastrointestinal tract. J. Dig. Dis. 21, 125–126.

Garcıá, S.R., Deprez, M., Lebrigand, K., Cavard, A., Paquet, A., Arguel, M.J., Magnone, V., Truchi, M., Caballero, I., Leroy, S., et al. (2019). Novel dynamics of human mucociliary differentiation revealed by single-cell RNA sequencing of nasal epithelial cultures. Dev.

Glowacka, I., Bertram, S., Muller, M.A., Allen, P., Soilleux, E., Pfefferle, S., Steffen, I., Tsegaye, T.S., He, Y., Gnirss, K., et al. (2011). Evidence that TMPRSS2 Activates the Severe Acute Respiratory Syndrome Coronavirus Spike Protein for Membrane Fusion and Reduces Viral Control by the Humoral Immune Response. J. Virol. 85, 4122–4134.

Gordon, D.E., Jang, G.M., Bouhaddou, M., Xu, J., Obernier, K., White, K.M., O’Meara, M.J., Rezelj, V. V., Guo, J.Z., Swaney, D.L., et al. (2020). A SARS-CoV-2 protein interaction map reveals targets for drug repurposing. Nature 1–13.

Gralinski, L.E., and Baric, R.S. (2015). Molecular pathology of emerging coronavirus infections. J. Pathol. 235, 185–195.

Gramberg, T., Hofmann, H., Möller, P., Lalor, P.F., Marzi, A., Geier, M., Krumbiegel, M., Winkler, T., Kirchhoff, F., Adams, D.H., et al. (2005). LSECtin interacts with filovirus glycoproteins and the spike protein of SARS coronavirus. Virology 340, 224–236.

Gu, J., Gong, E., Zhang, B., Zheng, J., Gao, Z., Zhong, Y., Zou, W., Zhan, J., Wang, S., Xie, Z., et al. (2005). Multiple organ infection and the pathogenesis of SARS. J. Exp. Med. 202, 415–424.

de Haan, C.A.M., and Rottier, P.J.M. (2005). Molecular Interactions in the Assembly of Coronaviruses. Adv. Virus Res. 64, 165–230.

Hamming, I., Timens, W., Bulthuis, M.L.C., Lely, A.T., Navis, G.J., and van Goor, H. (2004). Tissue distribution of ACE2 protein, the functional receptor for SARS coronavirus. A first step in understanding SARS pathogenesis. J. Pathol.

Han, X., Zhou, Z., Fei, L., Sun, H., Wang, R., Chen, Y., Chen, H., Wang, J., Tang, H., Ge, W., et al. (2020). Construction of a human cell landscape at single-cell level. Nature.

Hikmet, F., Mear, L., Uhlen, M., and Lindskog, C. (2020). The protein expression profile of ACE2 in human tissues. BioRxiv 16.

Hoffmann, M., Kleine-Weber, H., Schroeder, S., Krüger, N., Herrler, T., Erichsen, S., Schiergens, T.S., Herrler, G., Wu, N.H., Nitsche, A., et al. (2020). SARS-CoV-2 Cell Entry Depends on ACE2 and TMPRSS2 and Is Blocked by a Clinically Proven Protease Inhibitor. Cell 181, 271–280.e8.

Hosier, H., Farhadian, S., Morotti, R., Deshmukh, U., Lu-Culligan, A., Campbell, K., Yasumoto, Y., Vogels, C., Casanovas-Massana, A., Vijayakumar, P., et al. (2020). SARS-CoV-2 Infection of the Placenta. MedRxiv 2020.04.30.20083907.

Huang, C., Wang, Y., Li, X., and et al., (2020). Clinical features of patients infected with 2019 novel coronavirus in Wuhan, China. Lancet 395, 497–506.

Huang, I.-C., Bailey, C.C., Weyer, J.L., Radoshitzky, S.R., Becker, M.M., Chiang, J.J., Brass, A.L., Ahmed, A.A., Chi, X., Dong, L., et al. (2011). Distinct Patterns of IFITM-Mediated Restriction of Filoviruses, SARS Coronavirus, and Influenza A Virus. PLoS Pathog. 7, e1001258.

Izsvák, Z., Wang, J., Singh, M., Mager, D.L., and Hurst, L.D. (2016). Pluripotency and the endogenous retrovirus HERVH: Conflict or serendipity? BioEssays 38, 109–117.

Jia, H.P., Look, D.C., Shi, L., Hickey, M., Pewe, L., Netland, J., Farzan, M., Wohlford-Lenane, C., Perlman, S., and McCray, P.B. (2005). ACE2 Receptor Expression and Severe Acute Respiratory Syndrome Coronavirus Infection Depend on Differentiation of Human Airway Epithelia. J. Virol. 79, 14614–14621.

Jia, R., Pan, Q., Ding, S., Rong, L., Liu, S.-L., Geng, Y., Qiao, W., and Liang, C. (2012). The N-Terminal Region of IFITM3 Modulates Its Antiviral Activity by Regulating IFITM3 Cellular Localization. J. Virol. 86, 13697–13707.

John Hopkins University and Medicine (2020). COVID-19 Map - Johns Hopkins Coronavirus Resource Center.

Kam, Y.-W., Okumura, Y., Kido, H., Ng, L.F.P., Bruzzone, R., and Altmeyer, R. (2009). Cleavage of the SARS Coronavirus Spike Glycoprotein by Airway Proteases Enhances Virus Entry into Human Bronchial Epithelial Cells In Vitro. PLoS One 4, e7870.

Kong, S.L., Chui, P., Lim, B., and Salto-Tellez, M. (2009). Elucidating the molecular physiopathology of acute respiratory distress syndrome in severe acute respiratory syndrome patients. Virus Res. 145, 260–269.

Lamers, M.M., Beumer, J., van der Vaart, J., Knoops, K., Puschhof, J., Breugem, T.I., Ravelli, R.B.G., Paul van Schayck, J., Mykytyn, A.Z., Duimel, H.Q., et al. (2020). SARS-CoV-2 productively infects human gut enterocytes. Science (80-.). eabc1669.

Lee, J.J., Kopetz, S., Vilar, E., Shen, J.P., Chen, K., and Maitra, A. (2020). Relative Abundance of SARS-CoV-2 Entry Genes in the Enterocytes of the Lower Gastrointestinal Tract. BioRxiv.

Li, W., Moore, M.J., Vasllieva, N., Sui, J., Wong, S.K., Berne, M.A., Somasundaran, M., Sullivan, J.L., Luzuriaga, K., Greeneugh, T.C., et al. (2003). Angiotensin-converting enzyme 2 is a functional receptor for the SARS coronavirus. Nature 426, 450–454.

Liang, W., Feng, Z., Rao, S., Xiao, C., Xue, X., Lin, Z., Zhang, Q., and Qi, W. (2020). Diarrhoea may be underestimated: A missing link in 2019 novel coronavirus. Gut.

Ling, Y., Xu, S.B., Lin, Y.X., Tian, D., Zhu, Z.Q., Dai, F.H., Wu, F., Song, Z.G., Huang, W., Chen, J., et al. (2020). Persistence and clearance of viral RNA in 2019 novel coronavirus disease rehabilitation patients. Chin. Med. J. (Engl).

Litviňuková, M., Talavera-López, C., Maatz, H., Reichart, D., Worth, C.L., Lindberg, E.L., Kanda, M., Polanski, K., Fasouli, E.S., Samari, S., et al. (2020). Cells and gene expression programs in the adult human heart. BioRxiv 2020.04.03.024075.

Lu, R., Zhao, X., Li, J., Niu, P., Yang, B., Wu, H., Wang, W., Song, H., Huang, B., Zhu, N., et al. (2020). Genomic characterisation and epidemiology of 2019 novel coronavirus: implications for virus origins and receptor binding. Lancet.

Ma, L., Xie, W., Li, D., Shi, L., Mao, Y., Xiong, Y., Zhang, Y., and Zhang, M. (2020). Effect of SARS-CoV-2 infection upon male gonadal function: A single center-based study. MedRxiv 2020.03.21.20037267.

Mar, K.B., Rinkenberger, N.R., Boys, I.N., Eitson, J.L., McDougal, M.B., Richardson, R.B., and Schoggins, J.W. (2018). LY6E mediates an evolutionarily conserved enhancement of virus infection by targeting a late entry step. Nat. Commun. 9.

Marzi, A., Gramberg, T., Simmons, G., Möller, P., Rennekamp, A.J., Krumbiegel, M., Geier, M., Eisemann, J., Turza, N., Saunier, B., et al. (2004). DC-SIGN and DC-SIGNR Interact with the Glycoprotein of Marburg Virus and the S Protein of Severe Acute Respiratory Syndrome Coronavirus. J. Virol. 78, 12090–12095.

Matsuyama, S., Nagata, N., Shirato, K., Kawase, M., Takeda, M., and Taguchi, F. (2010). Efficient Activation of the Severe Acute Respiratory Syndrome Coronavirus Spike Protein by the Transmembrane Protease TMPRSS2. J. Virol. 84, 12658–12664.

Melin, A.D., Janiak, M.C., Marrone, F., Arora, P.S., and Higham, J.P. (2020). Comparative ACE2 variation and primate COVID-19 risk. BioRxiv 2020.04.09.034967.

Mille, J.K., and Whittaker, G.R. (2014). Host cell entry of Middle East respiratory syndrome coronavirus after two-step, furin-mediated activation of the spike protein. Proc. Natl. Acad. Sci. U. S. A. 111, 15214–15219.

Moriguchi, T., Harii, N., Goto, J., Harada, D., Sugawara, H., Takamino, J., Ueno, M., Sakata, H., Kondo, K., Myose, N., et al. (2020). A first case of meningitis/encephalitis associated with SARS-Coronavirus-2. Int. J. Infect. Dis. 94, 55–58.

Ng, D.L., Al Hosani, F., Keating, M.K., Gerber, S.I., Jones, T.L., Metcalfe, M.G., Tong, S., Tao, Y., Alami, N.N., Haynes, L.M., et al. (2016). Clinicopathologic, immunohistochemical, and ultrastructural findings of a fatal case of middle east respiratory syndrome coronavirus infection in the United Arab Emirates, April 2014. Am. J. Pathol.

Paules, C.I., Marston, H.D., and Fauci, A.S. (2020). Coronavirus Infections-More Than Just the Common Cold. JAMA - J. Am. Med. Assoc.

Pfaender, S., Mar, K.B., Michailidis, E., Kratzel, A., Hirt, D., V’kovski, P., Fan, W., Ebert, N., Stalder, H., Kleine-Weber, H., et al. (2020). LY6E impairs coronavirus fusion and confers immune control of viral disease. BioRxiv 2020.03.05.979260.

Prasanth, K.R., Hirano, M., Fagg, W.S., McAnarney, E.T., Shan, C., Xie, X., Hage, A., Pietzsch, C.A., Bukreyev, A., Rajsbaum, R., et al. (2020). Topoisomerase III-ß is required for efficient replication of positive-sense RNA viruses. BioRxiv 2020.03.24.005900.

Qi, D., Yan, X., Tang, X., Peng, J., Yu, Q., Feng, L., Yuan, G., Zhang, A., Chen, Y., Yuan, J., et al. (2020). Epidemiological and clinical features of 2019-nCoV acute respiratory disease cases in Chongqing municipality, China: a retrospective, descriptive, multiple-center study. MedRxiv 2020.03.01.20029397.

Rozenblatt-Rosen, O., Stubbington, M.J.T., Regev, A., and Teichmann, S.A. (2017). The Human Cell Atlas: From vision to reality. Nature 550, 451–453.

Shi, S., Qin, M., Shen, B., Cai, Y., Liu, T., Yang, F., Gong, W., Liu, X., Liang, J., Zhao, Q., et al. (2020). Association of Cardiac Injury with Mortality in Hospitalized Patients with COVID-19 in Wuhan, China. JAMA Cardiol.

Simmons, G., Zmora, P., Gierer, S., Heurich, A., and Pöhlmann, S. (2013a). Proteolytic activation of the SARS-coronavirus spike protein: Cutting enzymes at the cutting edge of antiviral research. Antiviral Res. 100, 605–614.

Simmons, G., Zmora, P., Gierer, S., Heurich, A., and Pöhlmann, S. (2013b). Proteolytic activation of the SARS-coronavirus spike protein: Cutting enzymes at the cutting edge of antiviral research. Antiviral Res. 100, 605–614.

Sohni, A., Tan, K., Song, H.W., Burow, D., de Rooij, D.G., Laurent, L., Hsieh, T.C., Rabah, R., Hammoud, S.S., Vicini, E., et al. (2019). The Neonatal and Adult Human Testis Defined at the Single-Cell Level. Cell Rep. 26, 1501–1517.e4.

Sungnak, W., Huang, N., Bécavin, C., Berg, M., Queen, R., Litvinukova, M., Talavera-López, C., Maatz, H., Reichart, D., Sampaziotis, F., et al. (2020). SARS-CoV-2 entry factors are highly expressed in nasal epithelial cells together with innate immune genes. Nat. Med.

Tan, Y.W., Hong, W., and Liu, D.X. (2012). Binding of the 5’-untranslated region of coronavirus RNA to zinc finger CCHC-type and RNA-binding motif 1 enhances viral replication and transcription. Nucleic Acids Res.

Tang, T., Bidon, M., Jaimes, J.A., Whittaker, G.R., and Daniel, S. (2020). Coronavirus membrane fusion mechanism offers a potential target for antiviral development. Antiviral Res. 178, 104792.

Travaglini, K.J., Nabhan, A.N., Penland, L., Sinha, R., Gillich, A., Sit, R. V, Chang, S., Conley, S.D., Mori, Y., Seita, J., et al. (2019). A molecular cell atlas of the human lung from single cell RNA sequencing. BioRxiv 7191, 742320.

Vankadari, N., and Wilce, J.A. (2020). Emerging WuHan (COVID-19) coronavirus: glycan shield and structure prediction of spike glycoprotein and its interaction with human CD26. Emerg. Microbes Infect. 9, 601–604.

Venkatakrishnan, A., Puranik, A., Anand, A., Zemmour, D., Yao, X., Wu, X., Chilaka, R., Murakowski, D.K., Standish, K., Raghunathan, B., et al. (2020). Knowledge synthesis from 100 million biomedical documents augments the deep expression profiling of coronavirus receptors. BioRxiv 2020.03.24.005702.

Vento-Tormo, R., Efremova, M., Botting, R.A., Turco, M.Y., Vento-Tormo, M., Meyer, K.B., Park, J.E., Stephenson, E., Polański, K., Goncalves, A., et al. (2018). Single-cell reconstruction of the early maternal–fetal interface in humans. Nature 563, 347–353.

Vieira Braga, F.A., Kar, G., Berg, M., Carpaij, O.A., Polanski, K., Simon, L.M., Brouwer, S., Gomes, T., Hesse, L., Jiang, J., et al. (2019). A cellular census of human lungs identifies novel cell states in health and in asthma. Nat. Med.

Volunteers, A.-2019-nCoV, Li, Z., Wu, M., Guo, J., Yao, J., Liao, X., Song, S., Han, M., Li, J., Duan, G., et al. (2020). Caution on Kidney Dysfunctions of 2019-nCoV Patients. MedRxiv 2020.02.08.20021212.

Wagner, M., Yoshihara, M., Douagi, I., Damdimopoulos, A., Panula, S., Petropoulos, S., Lu, H., Pettersson, K., Palm, K., Katayama, S., et al. (2020). Single-cell analysis of human ovarian cortex identifies distinct cell populations but no oogonial stem cells. Nat. Commun. 11.

Walls, A.C., Park, Y.J., Tortorici, M.A., Wall, A., McGuire, A.T., and Veesler, D. (2020). Structure, Function, and Antigenicity of the SARS-CoV-2 Spike Glycoprotein. Cell 181, 281–292.e6.

Wang, Z., and Xu, X. (2020). scRNA-seq Profiling of Human Testes Reveals the Presence of the ACE2 Receptor, A Target for SARS-CoV-2 Infection in Spermatogonia, Leydig and Sertoli Cells. Cells 9, 920.

Wang, D., Hu, B., Hu, C., Zhu, F., Liu, X., Zhang, J., Wang, B., Xiang, H., Cheng, Z., Xiong, Y., et al. (2020a). Clinical Characteristics of 138 Hospitalized Patients with 2019 Novel Coronavirus-Infected Pneumonia in Wuhan, China. JAMA - J. Am. Med. Assoc. 323, 1061–1069.

Wang, K., Chen, W., Zhou, Y.-S., Lian, J.-Q., Zhang, Z., Du, P., Gong, L., Zhang, Y., Cui, H.-Y., Geng, J.-J., et al. (2020b). SARS-CoV-2 invades host cells via a novel route: CD147-spike protein. BioRxiv 2020.03.14.988345.

Wang, S., Zhou, X., Zhang, T., and Wang, Z. (2020c). The need for urogenital tract monitoring in COVID-19. Nat. Rev. Urol.

Wang, W., Xu, Y., Gao, R., Lu, R., Han, K., Wu, G., and Tan, W. (2020d). Detection of SARS-CoV-2 in Different Types of Clinical Specimens. JAMA - J. Am. Med. Assoc.

Wolock, S.L., Lopez, R., and Klein, A.M. (2019). Scrublet: Computational Identification of Cell Doublets in Single-Cell Transcriptomic Data. Cell Syst. 8, 281–291.e9.

Wrapp, D., Wang, N., Corbett, K.S., Goldsmith, J.A., Hsieh, C.L., Abiona, O., Graham, B.S., and McLellan, J.S. (2020). Cryo-EM structure of the 2019-nCoV spike in the prefusion conformation. Science (80-.).

Wu, Y., Guo, C., Tang, L., Hong, Z., Zhou, J., Dong, X., Yin, H., Xiao, Q., Tang, Y., Qu, X., et al. (2020a). Prolonged presence of SARS-CoV-2 viral RNA in faecal samples. Lancet Gastroenterol. Hepatol. 5, 434–435.

Wu, Y., Xu, X., Chen, Z., Duan, J., Hashimoto, K., Yang, L., Liu, C., and Yang, C. (2020b). Nervous system involvement after infection with COVID-19 and other coronaviruses. Brain. Behav. Immun.

Wu, C., Zheng, S., Chen, Y., and Zheng, M. (2020). Single-cell RNA expression profiling of ACE2, the putative receptor of Wuhan 2019-nCoV, in the nasal tissue. MedRxiv 2020.02.11.20022228.

Wyler, E., Mösbauer, K., Franke, V., Diag, A., Gottula, L.T., Arsie, R., Klironomos, F., Koppstein, D., Ayoub, S., Buccitelli, C., et al. (2020). Bulk and single-cell gene expression profiling of SARS-CoV-2 infected human cell lines identifies molecular targets for therapeutic intervention. BioRxiv 2020.05.05.079194.

Xiao, F., Tang, M., Zheng, X., Liu, Y., Li, X., and Shan, H. (2020). Evidence for Gastrointestinal Infection of SARS-CoV-2. Gastroenterology 158, 1831–1833.e3.

Xu, S., and Li, Y. (2020). Beware of the second wave of COVID-19. Lancet.

Xu, J., Qi, L., Chi, X., Yang, J., Wei, X., Gong, E., Peh, S., and Gu, J. (2006). Orchitis: A Complication of Severe Acute Respiratory Syndrome (SARS)1. Biol. Reprod. 74, 410– 416.

Xu, Y., Li, X., Zhu, B., Liang, H., Fang, C., Gong, Y., Guo, Q., Sun, X., Zhao, D., Shen, J., et al. (2020a). Characteristics of pediatric SARS-CoV-2 infection and potential evidence for persistent fecal viral shedding. Nat. Med. 26, 502–505.

Xu, Z., Shi, L., Wang, Y., Zhang, J., Huang, L., Zhang, C., Liu, S., Zhao, P., Liu, H., Zhu, L., et al. (2020b). Pathological findings of COVID-19 associated with acute respiratory distress syndrome. Lancet Respir. Med. 8, 420–422.

Yan, L., Yang, M., Guo, H., Yang, L., Wu, J., Li, R., Liu, P., Lian, Y., Zheng, X., Yan, J., et al. (2013). Single-cell RNA-Seq profiling of human preimplantation embryos and embryonic stem cells. Nat. Struct. Mol. Biol. 20, 1131–1139.

Yang, Z.-Y., Huang, Y., Ganesh, L., Leung, K., Kong, W.-P., Schwartz, O., Subbarao, K., and Nabel, G.J. (2004). pH-Dependent Entry of Severe Acute Respiratory Syndrome Coronavirus Is Mediated by the Spike Glycoprotein and Enhanced by Dendritic Cell Transfer through DC-SIGN. J. Virol. 78, 5642–5650.

Yeager, C.L., Ashmun, R.A., Williams, R.K., Cardellichio, C.B., Shapiro, L.H., Look, A.T., and Holmes, K. V. (1992). Human aminopeptidase N is a receptor for human coronavirus 229E. Nature 357, 420–422.

Young, M.D., and Behjati, S. (2018). SoupX removes ambient RNA contamination from droplet based single cell RNA sequencing data. BioRxiv 303727.

Young, B.E., Ong, S.W.X., Kalimuddin, S., Low, J.G., Tan, S.Y., Loh, J., Ng, O.T., Marimuthu, K., Ang, L.W., Mak, T.M., et al. (2020). Epidemiologic Features and Clinical Course of Patients Infected with SARS-CoV-2 in Singapore. JAMA - J. Am. Med. Assoc. 323, 1488–1494.

Yu, X.J., Luo, C., Lin, J.C., Hao, P., He, Y.Y., Guo, Z.M., Qin, L., Su, J., Liu, B.S., Huang, Y., et al. (2003). Putative hAPN receptor binding sites in SARS_CoV spike protein. Acta Pharmacol. Sin.

Zang, R., Castro, M.F.G., McCune, B.T., Zeng, Q., Rothlauf, P.W., Sonnek, N.M., Liu, Z., Brulois, K.F., Wang, X., Greenberg, H.B., et al. (2020). TMPRSS2 and TMPRSS4 mediate SARS-CoV-2 infection of human small intestinal enterocytes. BioRxiv 2020.04.21.054015.

Zeng, L., Xia, S., Yuan, W., Yan, K., Xiao, F., Shao, J., and Zhou, W. (2020). Neonatal Early-Onset Infection with SARS-CoV-2 in 33 Neonates Born to Mothers with COVID-19 in Wuhan, China. JAMA Pediatr.

Zhang, C., Shi, L., and Wang, F.S. (2020). Liver injury in COVID-19: management and challenges. Lancet Gastroenterol. Hepatol. 5, 428–430.

Zhao, Y., Zhao, Z., Wang, Y., Zhou, Y., Ma, Y., and Zuo, W. (2020). Single-cell RNA expression profiling of ACE2, the putative receptor of Wuhan 2019-nCov. BioRxiv.

Zhou, F., Yu, T., Du, R., Fan, G., Liu, Y., Liu, Z., Xiang, J., Wang, Y., Song, B., Gu, X., et al. (2020). Clinical course and risk factors for mortality of adult inpatients with COVID-19 in Wuhan, China: a retrospective cohort study. Lancet.

Ziegler, C., Allon, S.J., Nyquist, S.K., Mbano, I., Miao, V.N., Cao, Y., Yousif, A.S., Bals, J., Hauser, B.M., Feldman, J., et al. (2020). SARS-CoV-2 Receptor ACE2 is an Interferon-Stimulated Gene in Human Airway Epithelial Cells and Is Enriched in Specific Cell Subsets Across Tissues. SSRN Electron. J.

Zmora, P., Hoffmann, M., Kollmus, H., Moldenhauer, A.S., Danov, O., Braun, A., Winkler, M., Schughart, K., and Stefan Pöhlmann, X. (2018). TMPRSS11A activates the influenza A virus hemagglutinin and the MERS coronavirus spike protein and is insensitive against blockade by HAI-1. J. Biol. Chem. 293, 13863–13873.

Zou, L., Ruan, F., Huang, M., Liang, L., Huang, H., Hong, Z., Yu, J., Kang, M., Song, Y., Xia, J., et al. (2020a). SARS-CoV-2 viral load in upper respiratory specimens of infected patients. N. Engl. J. Med. 382, 1177–1179.

Zou, X., Chen, K., Zou, J., Han, P., Hao, J., and Han, Z. (2020b). Single-cell RNA-seq data analysis on the receptor ACE2 expression reveals the potential risk of different human organs vulnerable to 2019-nCoV infection. Front. Med.

